# Sexually dimorphic placental responses to maternal SARS-CoV-2 infection

**DOI:** 10.1101/2021.03.29.437516

**Authors:** Evan A Bordt, Lydia L Shook, Caroline Atyeo, Krista M Pullen, Rose M De Guzman, Marie-Charlotte Meinsohn, Maeva Chauvin, Stephanie Fischinger, Laura J. Yockey, Kaitlyn James, Rosiane Lima, Lael M Yonker, Alessio Fasano, Sara Brigida, Lisa M Bebell, Drucilla J Roberts, David Pépin, Jun R Huh, Staci D Bilbo, Jonathan Z Li, Anjali Kaimal, Danny Schust, Kathryn J Gray, Douglas Lauffenburger, Galit Alter, Andrea G Edlow

**Affiliations:** Department of Pediatrics, Lurie Center for Autism, Massachusetts General Hospital, Harvard Medical School, Boston, MA, USA.; Department of Obstetrics and Gynecology, Massachusetts General Hospital, Harvard Medical School, Boston, MA, USA; Vincent Center for Reproductive Biology, Massachusetts General Hospital, Boston, MA, USA; Ragon Institute of MGH, MIT, and Harvard, Cambridge, MA, USA; PhD Program in Virology, Division of Medical Sciences, Harvard University, Boston, MA, USA; Department of Biological Engineering, Massachusetts Institute of Technology, Cambridge, MA, USA; Pediatric Surgical Research Laboratories, Department of Surgery, Massachusetts General Hospital, Harvard Medical School, Boston, MA 02114, USA; Department of Medicine, Massachusetts General Hospital, Harvard Medical School, Boston, MA, USA; Mucosal Immunology and Biology Research Center, Massachusetts General Hospital, Boston, MA; Department of Pediatrics, Massachusetts General Hospital, Boston, MA; Harvard Medical School, Boston, MA, USA.; European Biomedical Research Institute of Salerno (EBRIS), Salerno, Italy.; Department of Medicine, Massachusetts General Hospital and Harvard Medical School, Boston, MA USA; Department of Pathology, Massachusetts General Hospital, Harvard Medical School, Boston, MA, USA; Department of Immunology, Blavatnik Institute, Harvard Medical School, Boston, MA, USA; Evergrande Center for Immunologic Diseases, Harvard Medical School and Brigham and Women’s Hospital, Boston, MA, USA; Duke University, Department of Psychology and Neuroscience, Durham, NC, USA; Department of Medicine, Brigham and Women’s Hospital, Boston, MA, USA; Department of Obstetrics, Gynecology, and Women’s Health, University of Missouri, Columbia, MO, USA; Department of Obstetrics and Gynecology, Brigham and Women’s Hospital, Harvard Medical School, Boston, MA, USA

**Author notes:** Address correspondence to: Andrea G. Edlow, Vincent Center for Reproductive Biology, Massachusetts General Hospital, 55 Fruit Street, Thier Research Building, 903b. These authors contributed equally to this work (Lydia L Shook, Caroline Atyeo, Krista M Pullen).

## Abstract

There is a persistent male bias in the prevalence and severity of COVID-19 disease. Underlying mechanisms accounting for this sex difference remain incompletely understood. Interferon responses have been implicated as a modulator of disease in adults, and play a key role in the placental anti-viral response. Moreover, the interferon response has been shown to alter Fc-receptor expression, and therefore may impact placental antibody transfer. Here we examined the intersection of viral-induced placental interferon responses, maternal-fetal antibody transfer, and fetal sex. Placental interferon stimulated genes (ISGs), Fc-receptor expression, and SARS-CoV-2 antibody transfer were interrogated in 68 pregnancies. Sexually dimorphic placental expression of ISGs, interleukin-10, and Fc receptors was observed following maternal SARS-CoV-2 infection, with upregulation in males. Reduced maternal SARS-CoV-2-specific antibody titers and impaired placental antibody transfer were noted in pregnancies with a male fetus. These results demonstrate fetal sex-specific maternal and placental adaptive and innate immune responses to SARS-CoV-2.

## INTRODUCTION

Mortality and morbidity risk during the perinatal period and infancy is significantly higher in males than in females (*1–4*). The underlying susceptibility of males may relate to evolutionary differences that occur throughout pregnancy and in the perinatal period, but the precise mechanistic differences that lead to this differential female survival benefit is not completely understood. Consistent with perinatal male vulnerability in general, male infants and children fare significantly worse in the setting of SARS-CoV-2 infection, with higher rates of severe disease in infants and multisystem inflammatory syndrome in children (MIS-C), a SARS-CoV-2 inflammatory syndrome associated with shock, cardiac involvement, and significantly elevated markers of inflammation (*5–10*). The biological basis for the observed relative vulnerability of the male immune system to SARS-CoV-2 in pediatric populations is likely multifactorial (*11, 12*). Emerging data point to a retrograde impact of infant sex on maternal immunity (*13, 14*), with specific differences in innate immune signaling across fetal sex, which may contribute to a differential dialogue between female and male fetuses with their mothers. This differential dialogue may critically impact immunity across the dyad, leading to an alternative hypothesis underlying sex-based differences in perinatal life, including the possibility of differential transfer of immunity from the mother that may empower female and male infants with differential levels of immunity.

Type I and Type III interferons (IFN) are induced after innate recognition of viruses. Upon binding to their receptors, they induce expression of downstream effectors, interferon stimulated genes (ISGs), which inhibit viral infection by a number of different mechanisms (*15*). However, viruses have evolved to evade these IFN responses, and IFN responses can also be drivers of inflammatory pathology. Type I IFN signaling correlates strongly with pathogenicity and fatality in both SARS-CoV-1 and MERS-CoV infections (*16–18*). Dysregulated Type I IFN signaling is also associated with severe disease and drives pathogenicity in SARS-CoV-2 infection, in both humans and murine models (*17, 19–25*). Notably, sex differences in adult peripheral blood and pulmonary IFN signaling have been observed in both SARS-CoV-1 and SARS-CoV-2 infection (*26–28*), but there is a dearth of information about sex differences in fetal and pediatric populations. Type I and Type III IFN responses at the maternal-fetal interface play a crucial role in limiting viral infection but may also be drivers of abnormal development (*29–31*). Sex differences have been noted in the placental immune response to bacterial infection (*32, 33*) and to other prenatal alterations such as maternal stress and maternal high-fat diet (*12, 34, 35*). However, placental expression of interferon-stimulated genes (ISGs) in maternal SARS-CoV-2 infection, the role of sex differences, and potential impact on placental function have not yet been examined.

Newborn anti-viral immunity relies heavily on the placental transfer of maternal immunoglobulin-G (IgG) to the fetal circulation (*36–38*). Public health strategies to protect newborns from potentially devastating respiratory infections such as pertussis and influenza capitalize on the ability of the placenta to transfer vaccine-induced maternal IgG to the fetal circulation (*39, 40*). Fragment crystallizable (Fc) receptors in the placenta play an active role in antibody transfer from mother to fetus (*36, 37, 41–43*). FCγR1 expression relies on IFN, and emerging data have clearly demonstrated perturbed placental transfer in the setting of other coinfections including HIV (*44*) and malaria (*45*). Whether differences in inflammatory responses to SARS-CoV-2 infection could influence placental antibody transfer is unknown. In addition, little is known regarding sex differences in neonatal immune profiles and in maternal-fetal antibody transfer. Recent work has demonstrated reduced transplacental transfer of SARS-CoV-2-specific antibodies relative to influenza and pertussis antibodies (*46, 47*), and associated alterations in expression and localization of specific Fc receptors in the placenta (*47*), but sex differences in neonatal antibody-mediated immunity to SARS-CoV-2 and in placental receptors involved in antibody transfer have not yet been characterized.

Given known sex differences in immune responses to SARS-CoV-2 infection (*26*), the observed sex differences in disease prevalence and severity in the pediatric population including infants (*6–9*), and the placenta’s critical role as an immune organ mediating anti-viral responses and antibody transfer at the maternal-fetal interface (*31*), we sought to examine sex differences in the placental immune response to SARS-CoV-2 and how this may impact placental expression of receptors associated with antibody transfer. In a cohort of 38 participants infected with SARS-CoV-2 during pregnancy (19 male and 19 female fetuses) and a comparator group of 30 contemporaneously-enrolled pregnant women testing negative for SARS-CoV-2 (15 male and 15 female fetuses), striking differences were noted in placental transfer of antibodies to male and female infants linked to different ISG-driven Fc-receptor expression profiles in the placenta. These data point to unexpected sex-driven bilateral communication across the maternal:fetal interface associated with differences in rates of antibody transfer to SARS-CoV-2 that may help explain a major unaddressed sex-based difference across infectious diseases in male infants globally.

## RESULTS

### Demographic and clinical characteristics of study participants

Maternal demographic and clinical characteristics of study participants for placental analyses grouped by offspring sex are depicted in Table 1, while those providing maternal and umbilical cord blood are depicted in Supplementary Table 1. Of the 68 participants, 34 were pregnant with females and 34 with males. There were no significant differences between groups with respect to maternal age, parity, obesity, diabetes, hypertension, or gestational age at delivery, among other characteristics examined. Women with SARS-CoV-2 infection during pregnancy were more likely to be Hispanic compared to negative controls, concordant with our prior report of ethnic disparities in COVID-19 vulnerability in our hospital catchment area (*48*). Of the 38 women with SARS-CoV-2 infection during pregnancy, there were no significant differences between male and female fetuses with respect to gestational age at diagnosis of SARS-CoV-2 infection, days from positive SARS-CoV-2 test to delivery, or severity of COVID-19 illness. Although both male and female fetuses of mothers with SARS-CoV-2 had lower birthweights than their sex-matched control counterparts, clinically this difference was not meaningful as all neonatal birthweights were in the normal range. No neonates born to mothers with SARS-CoV-2 infection during pregnancy were infected with SARS-CoV-2.

**Table 1.** Demographic and clinical characteristics of study cohort by fetal sex and maternal SARS-CoV-2 status. Abbreviations: DM/GDM, diabetes mellitus/gestational diabetes mellitus; BMI, body mass index; GA, gestational age; N/A, not applicable. ^a^ SARS-CoV-2 positive and negative status determined by nasopharyngeal RT PCR at time of sample collection. If a participant was SARS-CoV-2 positive at any time in pregnancy, she was included in “SARS-CoV-2 positive” category. ^b^ Significant differences between groups were determined using chi-square test for categorical variables, and Kruskal-Wallis test for continuous variables presented as median [IQR]. ^c^ Severity determinations were made based on published criteria from the Society for Maternal-Fetal Medicine and the National Institutes of Health.

|  | All (68) | Female |  | Male |  | P-value<sup>b</sup> |
| --- | --- | --- | --- | --- | --- | --- |
|  |  | SARS-CoV-2 negative (15) | SARS-CoV-2 positive (19)<sup>a</sup> | SARS-CoV-2 negative (15) | SARS-CoV-2 positive (19)<sup>a</sup> |  |
| Maternal Age, years | 33 [28-38] | 33 [30-36] | 34 [30-40] | 30 [27-37] | 31 [26-36] | 0.34 |
| Parity, *n* | 1 [0-2] | 1 [0-2] | 1 [0-2] | 1 [0-2] | 1 [0-2] | 0.34 |
| Race, *n* (%) |  |  |  |  |  | 0.19 |
| *White* | 40 (59) | 11 (73) | 8 (42) | 12 (80) | 9 (59) |  |
| *Black* | 5 (7) | 1 (7) | 3 (16) | 0 (0) | 1 (7) |  |
| *Asian* | 1 (1) | 0 (0) | 0 (0) | 1 (7) | 0 (0) |  |
| *Other* | 12 (18) | 3 (20) | 4 (21) | 1 (7) | 4 (8) |  |
| *Not Reported* | 10 (15) | 0 (0) | 4 (21) | 1 (7) | 5 (15) |  |
| Ethnicity, *n* (%) |  |  |  |  |  | 0.01 |
| *Hispanic* | 32 (47) | 4 (27) | 11 (58) | 2 (13) | 15 (79) |  |
| *Non-Hispanic* | 33 (48) | 10 (67) | 7 (37) | 12 (80) | 4 (21) |  |
| *Not Reported* | 3 (4) | 1 (7) | 1 (5) | 1 (7) | 0 (0) |  |
| Chronic hypertension, *n* (%) | 0 (0) | 0 (0) | 0 (0) | 0 (0) | 0 (0) | N/A |
| DM/GDM, *n* (%) | 6 (9) | 1 (7) | 3 (16) | 1 (7) | 1 (5) | 0.67 |
| BMI ≥ 30 kg/m<sup>2</sup>, *n* (%) | 20 (29) | 2 (13) | 7 (37) | 3 (20) | 8 (42) | 0.21 |
| Pre-pregnancy BMI, kg/m<sup>2</sup> | 27.4 [21.5-30.8] | 25.8 [21.6-29.4] | 28.5 [26.5-32.7] | 21.5 [20.3-29.8] | 29.5 [22.3-32.0] | 0.06 |
| GA at delivery, weeks | 39.2 [38.6-40.3] | 39.3 [39.0-39.4] | 39.3 [38.4-40.1] | 39.1 [39.0-41.1] | 39.0 [35.3-40.3] | 0.09 |
| Any labor, *n* (%) | 52 (76) | 8 (53) | 16 (84) | 11 (73) | 17 (89) | 0.07 |
| Neonatal birthweight, *g* | 3283 [3005-3590] | 3400 [3060-3590] | 3255 [2920-3340] | 3435 [3260-3730] | 3115 [2615-3590] | 0.06 |
| GA at positive SARS-CoV-2 test, weeks | 38.7 [35.6-39.6] | N/A | 36.3 [32.4-39.4] | N/A | 36.3 [32.4-39.5] | 0.90 |
| COVID-19 disease severity at diagnosis<sup>c</sup>, *n* |  |  |  |  |  | 0.82 |
| *Asymptomatic* | 16 (42) | N/A | 8 (42) | N/A | 8 (42) |  |
| *Mild/Moderate* | 19 (50) | N/A | 9 (47) | N/A | 10 (53) |  |
| *Severe/Critical* | 3 (8) | N/A | 2 (11) | N/A | 1 (5) |  |
| Time between SARS-CoV-2 symptom onset and delivery, days | 36.5 [18-57] | N/A | 44 [28-64] | N/A | 32 [8-51] | 0.18 |

### Sexually dimorphic placental expression of interferon stimulated genes (ISGs) in response to maternal SARS-CoV-2 infection

Dysregulation of placental type I IFN responses has been noted in response to other viral infections (*30, 31, 49*). Placental type III IFN responses play an essential barrier function in maternal viral infection (*50*). Given the key role ISGs are known to play in the placental anti-viral response and previous reports of sex differences in expression of ISGs in the plasma of patients with COVID-19 (*26*), we examined whether maternal SARS-CoV-2 infection was associated with sex-specific alterations in placental ISG response (Fig. 1A). There was a sexually dimorphic expression pattern of the classical ISGs *IFI6, CXCL10,* and *OAS1* driven by increased expression in male SARS-CoV-2-exposed placentas compared to SARS-CoV-2-negative controls (Fig. 1B-D; Supplementary Table 2). We next examined expression of *CCL2/MCP-1*, a type I IFN-stimulated cytokine implicated in monocyte chemotaxis (*51*) and upregulated in lung samples from patients with COVID-19 (*23, 24*), as well as *MX1*, an anti-viral response gene induced by type I/III IFNs and upregulated during SARS-CoV-2 infection (*52*). Maternal SARS-CoV-2 exposure was associated with increased expression of *CCL2* and *MX1* in the placenta, with these differences primarily driven by increases in male placentas (Fig. 1E-F; Supplementary Table 2). Although maternal SARS-CoV-2 infection did not affect expression of placental *TNF, IL6,* or *CCL7* compared to expression in placentas from women negative for SARS-CoV-2 (Supplementary Fig. 1A-C; Supplementary Table 2), there was a sexually dimorphic effect of maternal SARS-CoV-2 infection on the expression of anti-inflammatory factor *IL10* (*53*), driven by significantly increased expression in SARS-CoV-2-exposed male placentas (Fig. 1G; Supplementary Table 2). We also observed a male-specific increase in density of CD163+ Hofbauer cells, placental resident fetal macrophages (*54*), in response to maternal SARS-CoV-2 exposure (Figure 1H-I; Supplementary Table 2). While placental Hofbauer cell hyperplasia has been described in maternal SARS-CoV-2 infection (*86, 87*), sex differences have not yet been examined. Together, these results suggest a sex-specific, and in some cases, sexually dimorphic response of placental ISGs to maternal SARS-CoV-2 infection, with significant upregulation of ISGs in male placentas exposed to maternal SARS-CoV-2 infection.

**Figure 1.**
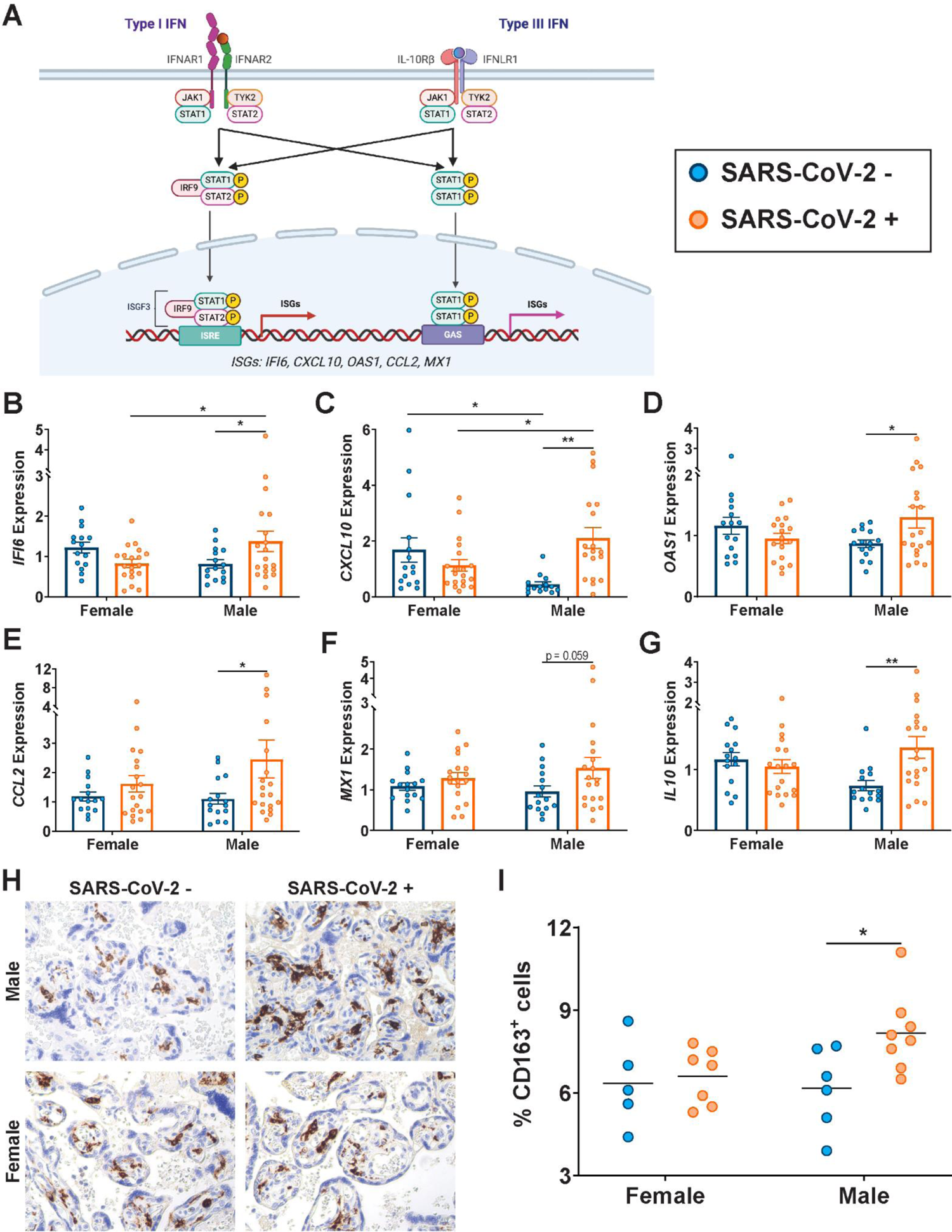
Male-specific upregulation of interferon stimulated genes in placentas exposed to maternal SARS-CoV-2 infection. **(A)** Interferon stimulated gene pathway diagram. Production of interferon stimulated genes (ISGs) of interest can occur through activation of Type I IFN (left) or Type III IFN (right). Image created using Biorender. **(B-G)** RTqPCR analyses of male or female placental expression of (B) *IFI6,* (C) *CXCL10,* (D) *OAS1*, (E) *CCL2*, (F) *MX1*, and (G) *IL10* in placental biopsies from SARS-CoV-2 negative (blue) or SARS-CoV-2 positive (orange) pregnancies. Expression levels shown are relative to reference gene *YWHAZ.* **(H-I)** Representative immunohistochemistry images and quantification of CD163-positive cells in placental slices from SARS-CoV-2 negative (blue) or SARS-CoV-2 positive (orange) pregnancies. Two-way ANOVA followed by Bonferroni’s post-hoc analyses were performed to determine significance. * p<0.05, ** p < 0.01

### Sex differences in placental transfer of SARS-CoV-2-specific antibodies and antibody function

Placental inflammation is associated with impaired antibody transfer (*45, 55*), and reduced transplacental antibody transfer has been shown in the context of maternal infections such as HIV (*44*) and malaria (*45*). Given the known vulnerability of male infants to more severe respiratory disease (*11*), we sought to determine whether the sex of the fetus would impact transplacental transfer of maternal SARS-CoV-2-specific antibodies. We comprehensively profiled SARS-CoV-2 antibodies spike (S), receptor binding domain (RBD), spike S1 subunit (S1), spike S2 subunit (S2), and nucleocapsid (N) including titer, function, and placental receptor interactions in maternal:cord plasma pairs of SARS-CoV-2 exposed and unexposed (SARS-CoV-2 negative) pregnancies, using a previously-described Systems Serology approach (*47*). Rather than the expected transplacental transfer observed for other pathogens (cord-to-maternal ratio of 1.1-1.5 or greater (*42, 56, 57*)), IgG titers against all examined SARS-CoV-2 antigens were significantly reduced in cord relative to maternal plasma across IgG subtypes in pregnancies with a male fetus compared to those with a female fetus (Fig. 2A, Supplementary Fig. 2). In contrast, significant reduction in cord versus maternal titers was only observed in the case of N-specific IgG2 in pregnancies with a female fetus (Supplementary Fig. 2). There were significantly reduced cord:maternal transfer ratios for SARS-CoV-2-specific antibodies in male versus female fetuses (Fig. 2B), and significantly reduced transfer of SARS-CoV-2-specific antibodies capable of binding to FcRn, FCGR2A/B and FCGR3A/B predominantly in males (Fig. 2C and Supplementary Fig. 2). Although transfer ratios for SARS-CoV-2-specific antibodies were significantly higher for female fetuses than those of males, they were still lower than the expected 1.1-1.5 ratios often observed for other pathogens (*47, 56, 57*). SARS-CoV-2-specific functional antibodies were also transferred less efficiently, with a marked male-specific decrease in antibody-dependent complement deposition (ADCD)-inducing antibodies and antibodies that induce natural killer (NK) cell chemokine secretion (macrophage inflammatory protein-1b/MIP-1b)(Fig. 2E-F). In contrast to SARS-CoV-2-specific antibodies, efficient transfer of pertussis antigen pertactin (PTN) and the influenza hemagglutinin (HA) glycoprotein-specific titers (Fig. 2C-D, Supplementary Fig. 3) and functions (Fig. 2E-F) was observed across pregnancies with both female and male fetuses.

**Figure 2.**
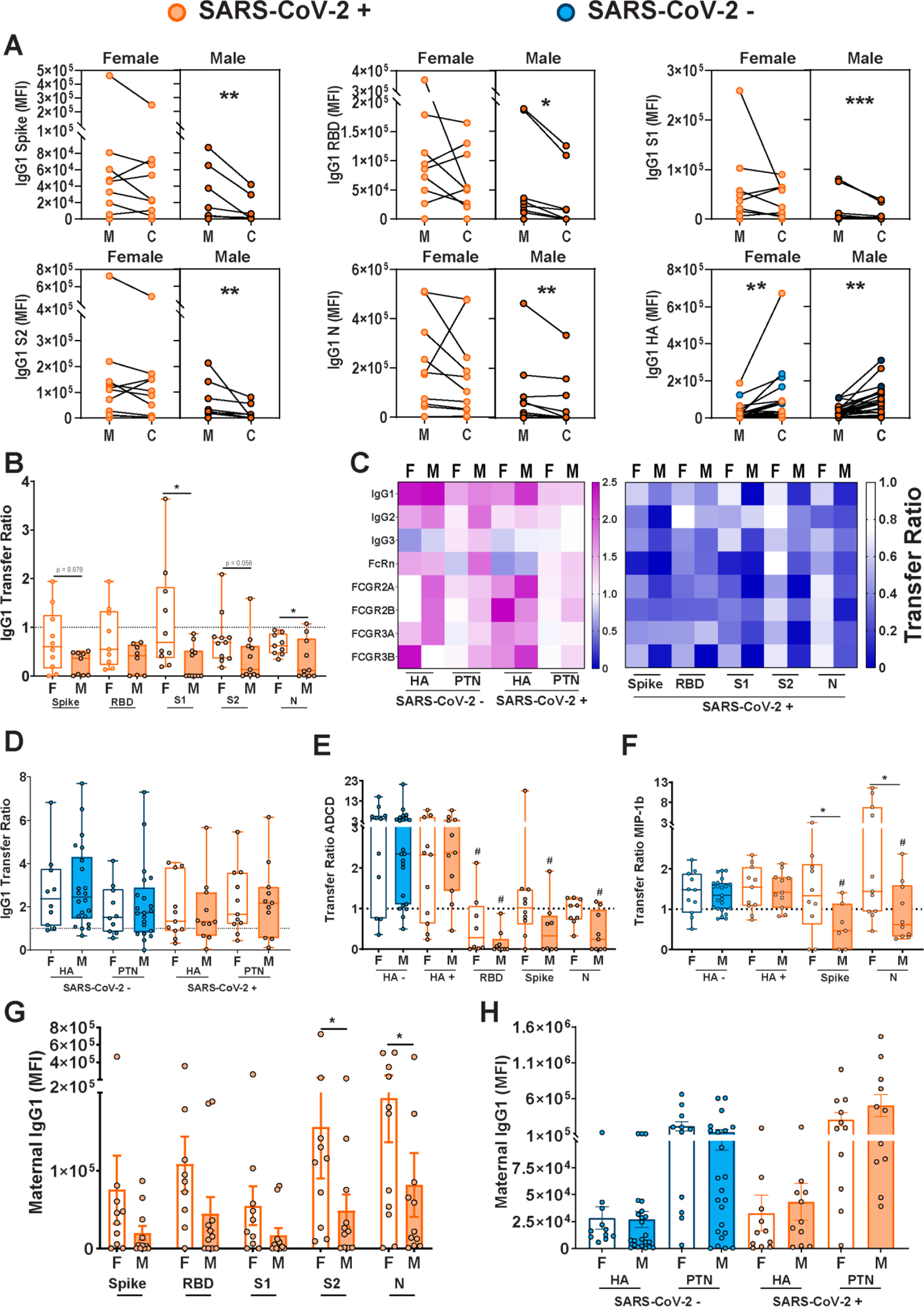
SARS-CoV-2 infected mothers with male fetuses demonstrate reduced placental transfer of SARS-CoV-2 antibodies compared to those with female fetuses. **A.** Dot plots showing relative Spike-, RBD-, S1-, S2-, N-, and influenza (HA)-specific maternal blood (M) and cord blood (C) titers of IgG1. Female neonates of SARS-CoV-2 negative mothers are shown in light blue, female neonates of SARS-CoV-2 positive mothers are shown in light orange. Males born to SARS-CoV-2 negative mothers are shown in dark blue, and males born to SARS-CoV-2 positive mothers are shown in dark orange. Units are Mean Fluorescence Intensity or MFI. All values reflect PBS background correction. Wilcoxon matched-pairs signed rank test was performed to determine significance. * p<0.05, ** p<0.01, *** p<0.001. **B.** Box-and-whisker plots show the transfer ratios (cord:maternal ratios) for IgG1 against the SARS-CoV-2 antigens Spike, RBD, S1, S2, and N. SARS-CoV-2 negative maternal status is shown in blue (female: open bars; male: shaded bars) and SARS-CoV-2 positive maternal status is shown in orange (female: open bars; male: shaded bars). All values are PBS background corrected. Kruskal-Wallis test followed by Dunn’s post-hoc analyses were performed to determine significance. In panels B-H, F indicates female and M indicates male fetus. Dotted line denotes a transfer ratio of 1 or 100%. * p<0.05. **C.** Heatmap depicting the median PBS background-corrected cord:maternal transfer ratio of HA, PTN, Spike, RBD, S1, S2, and N across all antibody subclasses and Fc receptor binding profiles in both SARS-CoV-2 negative and positive maternal:neonate dyads. **D.** The box-and-whisker plots show the transfer ratios (cord:maternal ratio) for IgG1 against HA and PTN in maternal:neonate dyads from either SARS-CoV-2 negative or SARS-CoV-2 positive pregnancies. SARS-CoV-2 negative are shown in blue (female: open bars; male: shaded bars) and SARS-CoV-2 positive are shown in orange (female: open bars; male: shaded bars). Kruskal-Wallis test followed by Dunn’s post-hoc analyses were performed to determine significance. In panels D-F, dotted line denotes transfer ratio of 1 or 100%. * p<0.05. **E-F.** Box-and-whisker plots depicting transfer ratios (cord:maternal) for (E) antibody-dependent complement deposition (ADCD) and (F) MIP-1b NK activation against HA, RBD, Spike, and N. SARS-CoV-2 negative maternal status is shown in blue (female: open bars; male: shaded bars) and SARS-CoV-2 positive maternal status is shown in orange (female: open bars; male: shaded bars). For each SARS-CoV-2 antigen, significance was determined against the HA-specific activity using the matched SARS-CoV-2 + dyad. Kruskal-Wallis test followed by post-hoc analyses was performed to determine significance. # p < 0.01 compared to HA SARS-CoV-2 + of that sex, * p < 0.01 compared to indicated group. **G.** Bar graphs depicting Spike-, RBD-, S1-, S2-, and N-specific maternal blood IgG1 levels. Two-way ANOVA followed by post-hoc analyses was used to determine significance. There was a main effect of fetal sex on maternal IgG1 levels. Units are MFI: Mean Fluorescence Intensity. *p < 0.05. **H.** Bar graphs depicting HA- and PTN-specific maternal blood IgG1 levels. SARS-CoV-2 negative maternal status is shown in blue (female: open bars; male: shaded bars) and SARS-CoV-2 positive maternal status is shown in orange (female: open bars; male: shaded bars). Two-way ANOVA revealed no effect of fetal sex or maternal SARS-CoV-2 infection status. Units are MFI: Mean Fluorescence Intensity.

As transplacental passive humoral immune transfer from maternal to cord blood is dependent in part on maternal antibody titers (*58*), we assessed the influence of fetal sex on maternal anti-SARS-CoV-2 antibody titers. Maternal levels of SARS-CoV-2-specific IgG1 were significantly lower in mothers with a male fetus compared to a female fetus (Fig. 2G). This effect was specific to anti-SARS-CoV-2 antibodies, with no significant impact of fetal sex on maternal anti-HA and PTN titers (Fig. 2H). Similar results were observed for maternal IgG2, IgG3, and IgM (Supplementary Fig. 4). Overall, these results suggest a reduced maternal SARS-CoV-2-specific humoral immune response in pregnancies with male fetuses. The reduced maternal antibody titers in mothers with male fetuses are consistent with prior studies demonstrating suppressed maternal pro-inflammatory responses in mothers with a male fetus relative to female fetus (*13, 14*) and the known direct correlation between pro-inflammatory response and increased antibody production in COVID-19 infection (*59, 60*).

### Sexually dimorphic placental Fc receptor expression in response to maternal SARS-CoV-2 infection

The transfer of maternal antibodies across the placenta is mediated by Fc receptors (*36, 37*). The neonatal Fc receptor (FcRn) was considered the classical receptor mediating transplacental transfer of IgG (*61*) (*62*), although other Fc receptors (FcγI,II,III) are increasingly recognized as being expressed in the placenta and likely to play a role in transfer (*36, 37*). Given the lower maternal antibody levels in pregnancies with a male fetus, we next aimed to probe the potential contribution of Fc-receptor expression to antibody-transfer differences across fetal sex. We observed sexually dimorphic expression of *FCGRT, FCGR1, FCGR3A* and *FCGR3B* in the setting of maternal SARS-CoV-2 infection (Fig 3A-C, Supplementary Fig.5 Supplementary Table 3), driven by increased expression in male SARS-CoV-2-exposed placentas. There were no fetal sex or maternal SARS-CoV-2 infection-mediated differences in placental expression of *FCGR2A/B* (Supplementary Fig. 5; Supplementary Table 3). These findings point to sexual dimorphism in Fc-receptor expression specifically in male infants, potentially in an effort to increase the transfer of antibodies to infants to empower them with enhanced immunity. To distinguish whether increased transcript expression of Fc receptors was compensatory in the setting of diminished protein expression, or whether protein expression mirrored gene expression, we performed immunoblot analyses of FcRn, FCγR3, FCγR2, and FCγR1. Protein expression was consistent with gene expression for FcRn, FCγR2 and FCγR3 (Fig. 3D-F, Supplementary Fig. 5 and Supplementary Table 3). There was no statistically significant impact of maternal SARS-CoV-2 infection nor fetal sex on protein expression of FCγR1 (Fig. 3E; Supplementary Table 3). In addition to Fc receptor quantity, it has been demonstrated that co-localization of other Fcγ receptors with FcRn augments efficiency of placental antibody transfer (*37, 47*). Immunohistochemical analyses of placental villi revealed significant sexual dimorphism in placental Fc receptor co-localization, with a significant increase in co-localization of FCγR3 and FcRn in male placentas only (Fig. 3G, 3J; Supplementary Table 3). No fetal sex or maternal SARS-CoV-2 exposure differences in co-localization with FcRn were observed for FCγR1 (Fig. 3H, 3K; Supplementary Table 3) or FCγR2 (Supplementary Fig. 5E-F; Supplementary Table 3). The gene and protein expression results suggest that maternal SARS-CoV-2 infection has a sexually dimorphic impact on placental Fcγ receptor expression, driven by an increase in overall expression and in FCγR3/FcRn co-localization in male placentas. Such increases may be compensatory for lower maternal titers of IgG in the setting of a male fetus, but still were not able to restore normal placental transfer of humoral immunity to male fetuses.

**Figure 3.**
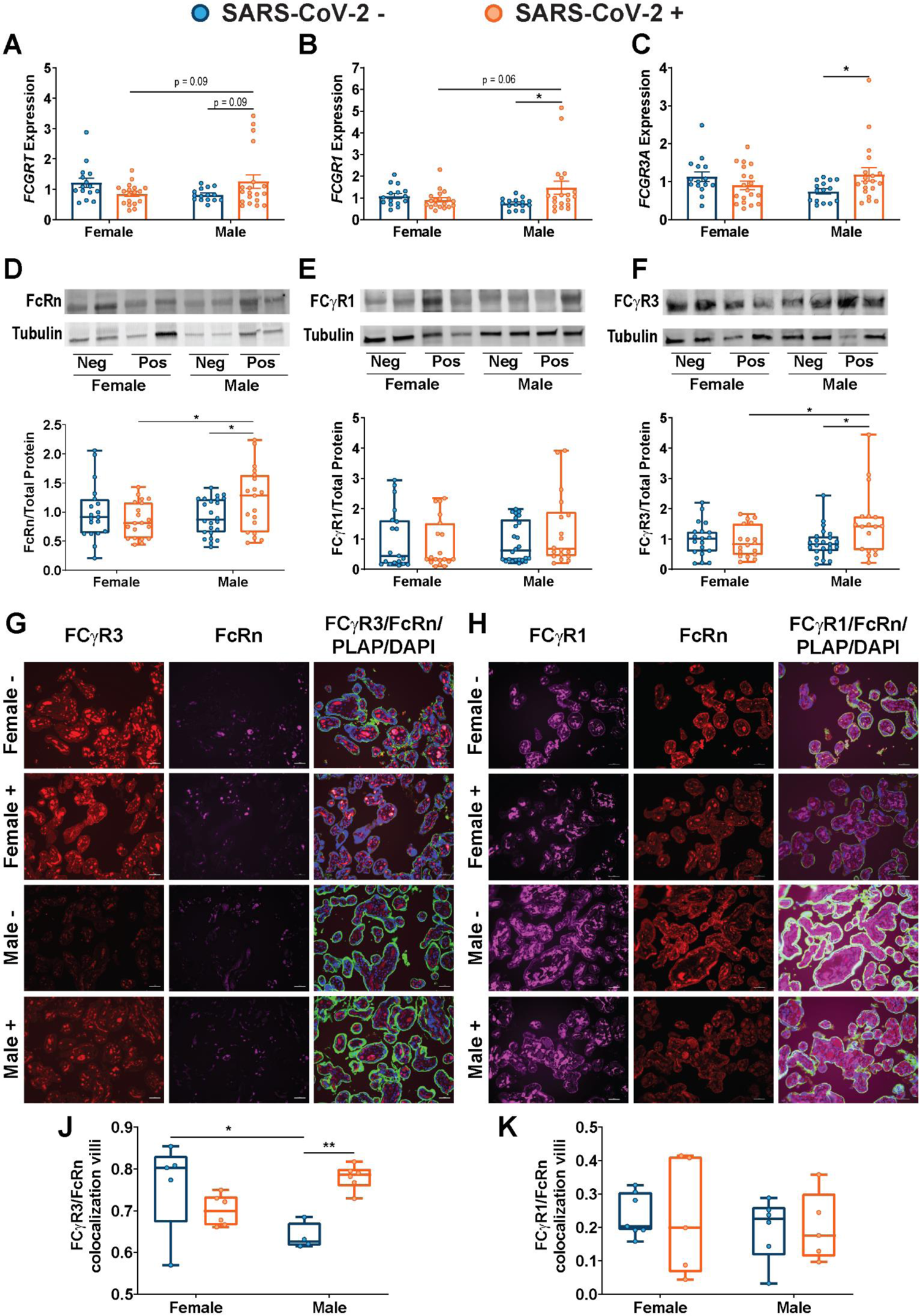
Sexually dimorphic regulation of Fc receptor gene expression, protein expression, and colocalization. **(A-C)** RTqPCR analyses of male or female placental expression of *FCGRT* (A)*, FCGR1* (B), and *FCGR3A* (C) in placental biopsies from SARS-CoV-2 negative (blue) or SARS-CoV-2 positive (orange) pregnancies. Expression levels shown are relative to reference gene *YWHAZ***. (D-F)** Representative immunoblots and quantification of fetal female or fetal male expression of FcRn (A), FCγR1 (B), and FCγR3 (C) in placental biopsies from SARS-CoV-2 negative (blue) or SARS-CoV-2 positive (orange) pregnancies. Neg and Pos on western blot designates SARS-CoV-2 negative and positive pregnancies. **(G)** Placental tissue sections from SARS-CoV-2 + and SARS-CoV-2 – mothers were stained for FCγR3 (red), FcRn (purple), and placental alkaline phosphatase (PLAP, green), a trophoblast marker, and DAPI (blue). **(H)** Box-and-whisker plots showing FCγR3/FcRn co-localization in placental villi. **(I)** Placental tissue sections from SARS-CoV-2 + and SARS-CoV-2 – mothers were stained for FCγR1 (purple), FcRn (red), and placental alkaline phosphatase (PLAP, green), a trophoblast marker, and DAPI (blue). **(J)** Box-and-whisker plots showing (J) FCγR1/FcRn or (K) FCγR2/FcRn co-localization in placental villi. Two-way ANOVA followed by Bonferroni’s post-hoc analyses were performed to determine significance. * p<0.05, ** p<0.01.

### Modeling the relative impact of placental gene expression, antibody features, and disease timing and severity

While we observed sexually dimorphic expression of individual placental genes and sex-biased transplacental antibody transfer in response to maternal SARS-CoV-2 infection, it was still unclear whether placental gene expression influenced the transfer of humoral immunity in a sex-specific manner. To investigate the relationship between fetal sex, placental gene expression, and antibody transfer, we examined whether S1-specific IgG1 transplacental transfer could be predicted by differential inflammatory, ISG, and Fc receptor placental gene expression in male and female orthogonal partial least squares discriminant analysis (O-PLSDA) models. S1-specific IgG1 transfer was modeled because the most significant sex difference was noted in the transfer of this antibody (Fig. 2B). Given that transfer of SARS-CoV-2 specific antibodies in fetuses of both sexes was less than the expected cord:maternal ratio of ≥1.1-1.5, there was no truly high or efficient transfer. Ratio thresholds for distinguishing higher and lower transfer were therefore determined based on the natural distribution of the data. For female neonates, this ratio was 0.8, and for males, this ratio was 0.4 (reflective of more impaired transfer in males). To investigate whether multivariate analysis could identify gene signatures specific to relatively high or low transfer, we performed O-PLSDA separately on the female and male neonates (Fig. 4A-D). To avoid overfitting the O-PLSDA model, features were reduced to the ‘optimal’ set of nonredundant features using LASSO feature selection (*63*). In both male and female pregnancies, high placental expression of *FCGR3B* was associated with lower IgG1 S1 transfer efficiency (Fig. 4). Interestingly, higher placental *IL10* expression was associated with higher IgG1 S1 transfer in female pregnancies, but lower transfer efficiency in male pregnancies (Fig. 4B,4D). In male pregnancies, high expression of *OAS1* was associated with higher transfer (Fig. 4D). Overall, these O-PLSDA models support a systemic sexually dimorphic response that triggers distinct inflammatory cascades likely critical for driving different levels of antibody production in mothers, as well as Fc-receptor expression alterations aimed at compensating for these differences.

**Figure 4.**
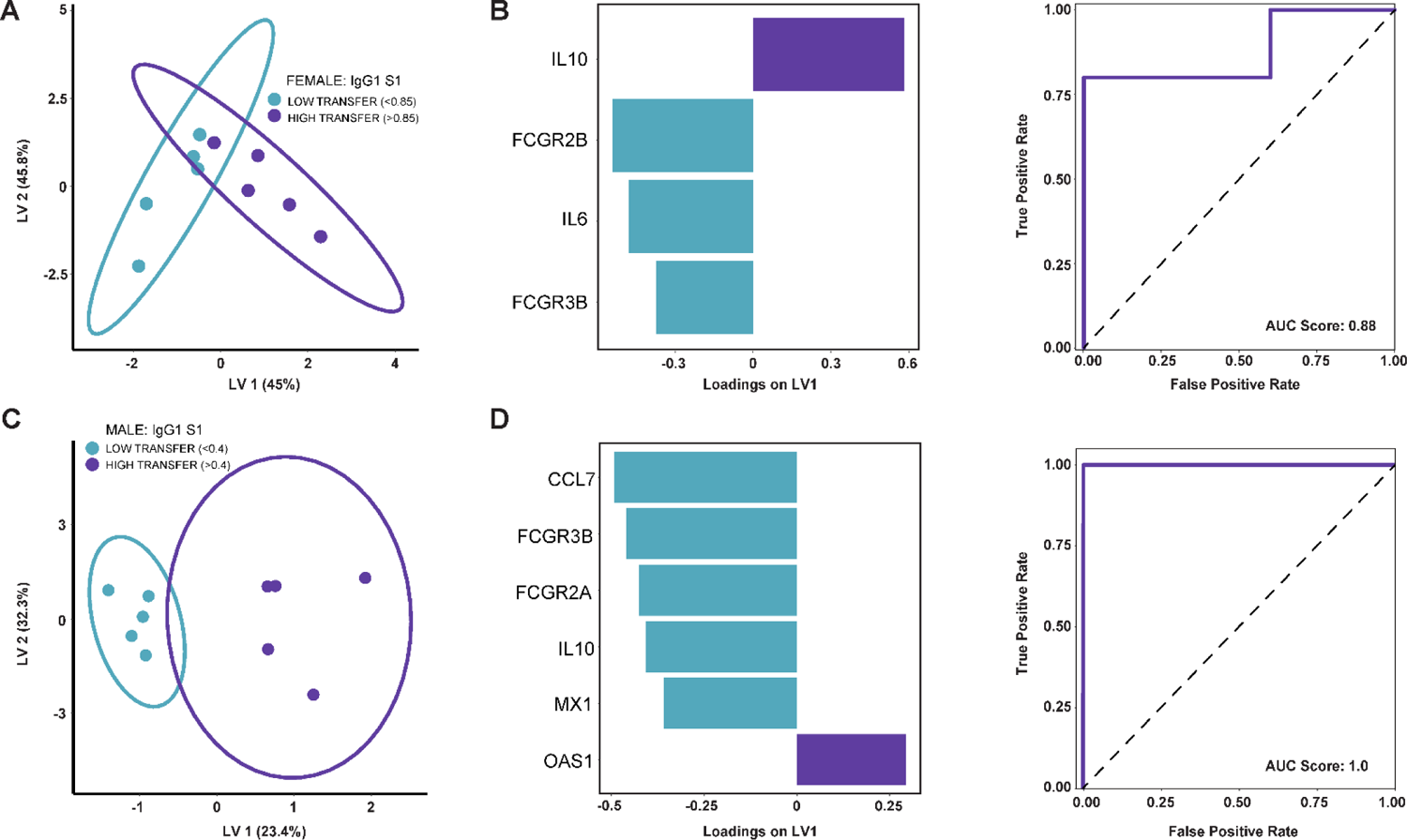
Transplacental antibody transfer characterized by gene expression. **(A)** OPLSDA model classifying relatively low (ratio < 0.8) vs. high (ratio > 0.8) IgG1 S1 transfer to female neonates based on placental gene expression. The model was built on gene expression features after LASSO feature selection. Each dot represents one mother-cord pair. **(B)** Gene expression feature loadings on the first latent variable (LV1) for the female model in (A). Bar color corresponds with which transfer classification (low vs. high) is characterized by enriched expression of that gene, with blue bars enriched in low transfer and purple bars enriched in high transfer. The receiver operating characteristic curve (solid purple line) for this model quantifies the accuracy in classifying IgG1 S1 transfer to female neonates after leave-one-out cross validation. The area under the curve (AUC) score is reported. The black dashed line represents the expected performance of a random model (AUC Score = 0.5). **(C)** OPLSDA model classifying relatively low (ratio < 0.4) vs. high (ratio > 0.4) IgG1 S1 transfer to male neonates based on placental gene expression. The model was built on gene expression features after LASSO feature selection. **(D)** Gene expression feature loadings on the first latent variable (LV1) for the male model in (C). Bar color corresponds with which transfer classification (low vs. high) is characterized by enriched expression of that gene, with blue bars enriched in low transfer and purple bars enriched in high transfer, respectively. The receiver operating characteristic curve (solid purple line) for this model quantifies the accuracy in classifying IgG1 S1 transfer to male neonates after leave-one-out cross validation. The AUC score is reported with black dashed line representing the expected performance of a random model (AUC Score = 0.5).

We also examined the sex-specific impacts of other disease and birth-related factors on placental ISG and Fc-receptor expression, and placental antibody transfer. Using linear regression we did not observe significant impact of time since infection or disease severity on male or female placental expression of any ISGs, pro-inflammatory cytokines, Fc receptors, or IgG transfer (Supplementary Fig. 6). Additionally, though both male and female neonates born to SARS-CoV-2 positive mothers had statistically significantly lower birthweight compared to sex-matched controls, birthweights were in the normal range for both groups (Table 1) and birthweight did not significantly impact IgG antibody transfer in linear regression models (Supplementary Fig 6). Similarly, we did not observe significant associations between antibody transfer and days from infection or gestational age at delivery for either male or female neonates (Supplementary Fig. 6). In summary, disease severity, time since infection, neonatal birthweight, and gestational age at delivery do not appear to confound our earlier findings related to sexually dimorphic gene expression and antibody transfer.

## DISCUSSION

Our results demonstrate the impact of fetal sex on the maternal and placental immune response to SARS-CoV-2, and the potential consequences for neonatal antibody-mediated immunity. We show that maternal SARS-CoV-2 infection is associated with sexually dimorphic expression of interferon stimulated genes and Fc receptors involved in placental transfer of maternal antibodies, with significant upregulation of ISGs and Fc receptors in SARS-CoV-2-exposed males. Male fetal sex was associated with reduced maternal SARS-CoV-2-specific IgG titers, and despite significant compensatory upregulation of placental Fc receptors, SARS-CoV-2-specific placental antibody transfer to the male fetus was reduced. Crosstalk between innate immunity (ISGs) and antibody adapters (FcRn/Fcγ) has been demonstrated to play a key role in optimizing tissue-specific immune responses against unfamiliar pathogens (*64, 65*). While Type I and III interferon responses may help protect the placenta from SARS-CoV-2 infection, the associated male-specific upregulation of placental Fc receptors was unable to overcome the deficit in SARS-CoV-2-specific antibody transfer from mother to fetus.

Epidemiologic data point to a persistent male bias in the development and severity of COVID-19 disease in adults, children, and infants (*6, 8, 9, 68, 69*). Male COVID-19 patients are three times as likely to require admission to intensive care units and have higher odds of death than females (*70*). This male-biased vulnerability to maternal SARS-CoV-2 infection is consistent with a male-biased risk of mortality and morbidity across the perinatal period in general (*1–3*). Our findings of sexually dimorphic placental innate immune responses to infection, coupled with sex differences in transfer of maternal humoral immunity, may reveal broader vulnerabilities contributing to higher morbidity and mortality in male infants.

Although the impact of fetal sex is not consistently evaluated in studies of placental function (*71*), sex-specific alterations in the placental transcriptome have been described in both normal and pathologic pregnancies (*72–75*). Sex differences in the placental immune response to prenatal infections and other immune stressors have been described in human and animal models (*32, 76–79*), but have not been examined in SARS-CoV-2 infection. Here we report that maternal SARS-CoV-2 infection induces a sexually dimorphic placental anti-viral innate immune response, with significant upregulation of ISGs in male, but not female, placentas. Male-specific stimulation of placental ISGs following SARS-CoV-2 exposure is consistent with the heightened male immune responses reported in SARS-CoV-2-infected adult and pediatric cohorts (*5, 6, 8–10, 26, 80*). Interestingly, although we do not see evidence of placental, maternal blood, or cord blood SARS-CoV-2 infection (*46*), and the majority of maternal infections represent mild or moderate disease, there is still evidence of altered placental gene expression and an anti-viral response in the placentas of male pregnancies. This indicates that even a mild maternal infection has the potential to affect the placenta and fetus.

Innate immune sensing of SARS-CoV-2 involves the activation of type I and type III interferons and upregulation of ISGs in target cells (*81*). Given the relative paucity of SARS-CoV-2 placental infection (*46*) in comparison to other pandemic infections such as Zika (*82*), the increased ISG production and upregulated *IL10* expression in exposed male placentas may be a protective mechanism to prevent direct placental infection and pathology. Indeed, high IFN levels during pregnancy have proven protective against placental HSV infection (*83*) and type III IFNs impair ZIKV transplacental transmission (*84*). Induction of ISGs is likely not universally protective, however. While type III IFNs primarily serve a barrier defense role, type I IFNs can serve a more classical immune activating/inflammatory role (*15, 29*). Animal models of viral infection in pregnancy implicate type I interferons and ISGs in impaired placental development and fetal growth restriction (*30, 85*), conditions which can have both short- and long-term impact on fetal and offspring health. Thus, it remains unclear if the male-specific upregulation of ISGs described here is potentially beneficial (e.g. protection from viral infection) or harmful, and what the long-term consequences may be for fetal development and in utero programming of later life metabolic and neurodevelopmental outcomes.

Due to their immature immune system, newborns rely on the passive transplacental transfer of maternal antibodies for initial protection against infectious pathogens (*36, 39, 86*). While previous reports in adults have noted sex differences in production of SARS-CoV-2-specific antibodies (*26, 87*), this is the first report of sex-biased maternal production and transplacental transfer of SARS-CoV-2-specific antibodies. We previously reported impaired placental transfer of maternal SARS-CoV-2-specific antibodies in the setting of maternal SARS-CoV-2 infection (*46, 47*), consistent with the reduced transplacental transfer of maternal antibodies in other maternal infections such as malaria and HIV (*38, 88–91*). While there are known sex differences in adult antibody production in response to SARS-CoV-2 infection (*26, 80*), little is known about sex differences in maternal titers or transplacental antibody transfer (*92, 93*), particularly in the setting of maternal SARS-CoV-2 infection. Our finding of impaired placental transfer of SARS-CoV-2-specific antibodies, more pronounced in males, is consistent with the male-specific reduction of placental transfer of maternal IgG reported in a non-human primate model of maternal stress (*92*). We observed decreased maternal antibody titers against all measured SARS-CoV-2-specific antigens (S, S1, S2, RBD, N) when the fetus was male compared to female, a difference not noted for influenza or pertussis antigens. The reduced maternal antibody titers in the setting of a male fetus are likely attributable to suppressed maternal pro-inflammatory responses in the setting of a male fetus, possibly to improve tolerance of the fetal allograft (*13, 14*), and the direct correlation between pro-inflammatory response and increased antibody production noted in COVID-19 infection (*59, 60*). Reduced maternal SARS-CoV-2-specific IgG titer in male pregnancies is undoubtedly a driver of reduced transplacental transfer noted in male fetuses, as maternal titer is a key aspect of passive antibody transfer (*94*). Whether the male-biased impairment in placental SARS-CoV-2-specific antibody transfer renders male infants more vulnerable to early-life SARS-CoV-2 infection remains unclear, as the amount of antibody necessary for protection against SARS-CoV-2 infection is unknown.

Differential induction of placental ISGs in males versus females may impact placental antibody transfer via alteration in Fc receptor expression (*95–97*) and function (*98*). FCγ receptor expression is upregulated on monocytes in the setting of Type I IFN signaling (*99*). The significant upregulation of *FCGRT, FCGR1,* and *FCGR3A/B* we observed in male SARS-CoV-2-exposed placentas may therefore be a response to significantly increased IFN signaling in exposed male placentas. Hofbauer cells, tissue-resident macrophages of the placenta, express FcγRI, II and III (*41*). The male-specific Hofbauer cell hyperplasia in placentas exposed to maternal SARS-CoV-2 infection could contribute to increased placental FCGR1/FCGR3 expression in males. Importantly, given the known role of placental Fc receptors in facilitating transplacental antibody transfer to the fetus, the male-specific upregulation of these receptors could be a compensatory response for the lower maternal SARS-CoV-2-specific antibody titers we describe in pregnancies with a male fetus.

A limitation of our study is the infection of participants primarily in the third trimester, because these samples were collected during the initial wave of the SARS-CoV-2 pandemic in Boston. Whether maternal SARS-CoV-2 infection in the first and second trimester alters ISG and Fc receptor expression, and how such altered expression might durably impact placental immune function, is a question that remains to be answered in future studies. In addition, although we found no significant association between disease severity and placental gene expression or antibody transfer, such examinations were limited by the relatively small number of women with severe or critical illness. Finally, while our regression models did not find time from infection to delivery to be a significant contributor to the antibody transfer ratios, we cannot entirely rule out any contribution of timing of maternal infection to the reduced antibody transfer noted in males. However, our robust sexually dimorphic gene and protein expression results, with significant upregulation of both placental ISGs and Fc receptors in males, demonstrate placental factors are a stronger driver of antibody transfer than any time-from-infection effect.

In conclusion, our comprehensive evaluation of the impact of fetal sex on placental gene expression and transplacental antibody transfer in maternal SARS-CoV-2 infection provides insight into sexually dimorphic or sex-specific placental innate and adaptive immune responses to maternal SARS-CoV-2 infection. The significantly increased impact of maternal SARS-CoV-2 infection on male placental and neonatal immunity highlights the importance of evaluating fetal sex in future studies of placental pathology and infant outcomes in SARS-CoV-2, as well as the critical importance of disaggregating sex data in follow-up studies of offspring neurodevelopmental and metabolic outcomes. These findings may have broader implications for understanding placental immune response, male vulnerability, and passive transfer of maternal antibody in other viral infections. Studies investigating SARS-CoV-2 vaccine safety and efficacy in pregnant women should also study placental immune response and antibody-transfer effects in addition to neonatal infection rates, and report these data in a sex-disaggregated fashion (*100*).

## METHODS

### Study design and participant recruitment

Pregnant women at two tertiary care centers in Boston, Massachusetts were approached for enrollment in a COVID-19 pregnancy biorepository study starting April 2, 2020 (Massachusetts General Hospital and Brigham and Women’s Hospital; Mass General Brigham IRB approval #2020P000804). All participants provided written informed consent (virtual). Pregnant women were eligible for inclusion if they were: (1) 18 years of age or older, (2) able to provide informed consent or had a healthcare proxy to do so, (3) diagnosed with, or at risk for, SARS-CoV-2 infection. Due to wide community spread in Massachusetts during the study period (*101*), all pregnant women presenting for hospital care were deemed “at risk” for SARS-CoV-2 infection. Enrollment coincided with universal COVID-19 screening of all patients admitted to the Labor and Delivery units by nasopharyngeal RT-PCR testing (initiated April 16, 2020). Maternal SARS-CoV-2 positivity was defined by a positive clinical nasopharyngeal RT-PCR test result.

### Maternal and cord blood collection and processing

Sample collection protocols have been described in a previous publication (*102*) and are described in detail in the Supplementary Materials and Methods.

#### Systems Serology

A multiplexed Luminex assay was used to determine relative titer of antigen-specific isotypes, subclasses, and FcR binding, as previously described (*45*). Detailed methodology is available in the Supplementary Materials and Methods.

### Placental sampling and processing

Fetal side biopsies (0.5 cubic centimeter per biopsy) were collected immediately after delivery. Biopsies were taken from two separate locations at least 4 cm from the cord insertion avoiding surface vasculature, edges, and areas of gross pathology. Fetal membranes were dissected from the fetal and maternal surfaces, respectively. Biopsies were serially washed in DPBS, submerged in 5 volumes of RNA*later*® (Invitrogen), and stored at least 24 hours at 4°C. Excess RNA*later*® was removed by blotting on Kimwipes, and biopsies were subdivided into 50 mg segments, placed into cryovials, flash frozen in liquid nitrogen and stored at −80°C. For histopathological examination, placentas were fixed in formalin, weighed, examined grossly, and sectioned/stained as described below.

### RNA extraction, cDNA synthesis, and qPCR

Placental tissue was placed in Trizol (100 uL/10 mg) and homogenized using a TissueTearor. Chloroform extraction was performed using an Rneasy Mini Kit (Qiagen), and cDNA synthesis was performed using iScript^TM^ cDNA Synthesis Kit (Bio-Rad) according to manufacturer instructions, as detailed in the Supplemental materials and methods.

### Protein extraction and quantification

Placental tissue was placed in RIPA Lysis and Extraction Buffer (ThermoFisher Scientific #89901) containing Halt Protease and Phosphatase Inhibitor Cocktail (ThermoFisher Scientific #78443) and homogenized using a TissueTearor. Resulting mixture was centrifuged at 10,000 rpm for 10 min at 4°C and supernatant was kept for protein quantification. Protein was quantified using Pierce BCA Protein Assay Kit (ThermoFisher Scientific #23225).

### Immunoblotting

50 ug of protein was prepared in Protein Sample Loading Buffer (Li-Cor Biosciences) and loaded on Mini-PROTEAN 4-20% TGX Stain-Free Precast Gels (Bio-Rad). Immunoblotting was performed as detailed in the Supplementary Materials and Methods. Fluorescence was then imaged with a ChemiDoc MP System (Bio-Rad) and quantified using Image Lab Software (Bio-Rad). Data is presented as volume of bands for protein of interest relative to the volume of bands contained in the total protein stain-free image.

### Immunofluorescence

For co-labeling immunofluoresence (IF) experiments, placenta tissue sections were prepared as previously described (*47*). Detailed methodology is available in the Supplementary Materials and Methods.

### Multivariate Analyses

All multivariate analyses were performed using R (version 4.0.0) and are detailed in the Supplementary Materials and Methods.

### Statistical Analyses

Statistical analyses were performed using GraphPad Prism 9 and using the ‘stats’ package in R (Version 4.0.0). Details of the specific statistical tests performed and statistical significance are described in figure legends and in the Supplementary Material.

## Author Contributions

Conceptualization and Methodology, E.A.B., L.L.S., G.A., A.G.E.; Investigation, E.A.B., L.L.S., C.A., K.M.P., R.M.D.G., M-C.M., M.C., S.F., L.J.Y., S.B., K.G., G.A., A.G.E; Formal Analysis, E.A.B., C.A., K.M.P., M-C.M., K.J., D.J.R., D.L., G.A., A.G.E.; Writing – Original Draft, E.A.B., L.L.S., C.A., G.A., A.G.E.; Writing-Review and Editing and Supervision, E.A.B., L.L.S., C.A., K.M.P., R.M.D.G., M-C.M., M.C., S.F., L.J.Y., K.J., R.L., L.M.Y., A.F., S.B., L.M.B., D.J.R., D.P., J.R.H., S.D.B., J.Z.L., A.K., D.J.S., K.J.G., D.L., G.A., and A.G.E.

## Data and materials availability

All data associated with this study are present in the paper or the Supplementary Materials.

## SUPPLEMENTARY MATERIALS

## METHODS

### Maternal and cord blood collection and processing

Sample collection protocols have been described in a previous publication (*102*) and are summarized below. Blood from pregnant women was collected at delivery by venipuncture into EDTA tubes. Umbilical cord blood was collected immediately after delivery. The umbilical cord was wiped clean and blood was drawn directly from the vein using a syringe and transferred to EDTA vacutainer tubes. Blood was centrifuged at 1000g for 10 min at room temperature. Plasma was aliquoted into cryogenic vials and stored at −80°C.

### Systems Serology

A multiplexed Luminex assay was used to determine relative titer of antigen-specific isotypes, subclasses, and FcR binding, as previously described (*45*). The following antigens were used in this assay: SARS-CoV-2 RBD (Sino Biological), SARS-CoV-2 Spike (LakePharma), SARS-CoV-2 N (Aalto Bio Reagents). SARS-CoV-2 S1 (Sino Biological), SARS-CoV-2 S2 (Sino Biological), pertussis pertactin (List Reagents) and a mix of HA A/Michigan/45/2015 (H1N1), HA A/Singapore/INFIMH-16-0019/2016 (H3N2), B/Phuket/3073/2013 (Immunetech). Antigens were covalently linked to carboxyl-modified Magplex© Luminex beads using Sulfo-NHS (N-hydroxysulfosuccinimide, Pierce) and ethyl dimethylaminopropyl carbodiimide hydrochloride (EDC). Antigen-coupled microspheres were blocked, washed, resuspended in PBS, and stored at 4°C. Plasma (diluted 1:100 for IgG2/3, 1:500 for IgG1 and FcRn, 1:1000 for all other FcRs) was added to the antigen-coupled microspheres to form immune complexes, and plates were incubated overnight at 4°C, shaking at 700 rpm. The next day, plates were washed with 0.1% BSA 0.02% Tween-20. Phycoerythrin (PE)-coupled mouse anti-human detection antibodies (Southern Biotech) were used to detect antigen-specific antibody binding. Avi-Tagged FcRs (Duke Human Vaccine Institute) were biotinylated using BirA500 kit (Avidity) per manufacturer’s instructions in order to detect FcR binding. Biotinylated FcRs were tagged with PE and added to immune complexes. Fluorescence was acquired using an Intellicyt iQue. Relative antigen-specific antibody titer and FcR binding is reported as Median Fluorescence Intensity (MFI).

### Placental sampling and processing

Fetal side biopsies (0.5 cubic centimeter per biopsy) were collected immediately after delivery from two separate locations at least 4 cm from the cord insertion avoiding surface vasculature, edges, and areas of gross pathology. Fetal membranes were dissected from the fetal surface and not included in the sample. Biopsies were then dissected in half, serially washed in DPBS and placed in RNA*later*® (Invitrogen) and stored at least 24 hours at 4°C. Excess RNA*later*® was then removed by blotting on Kimwipes, and biopsies were subdivided into 50mg segments, placed into cryovials, flash frozen in liquid nitrogen and stored at −80°C. For histopathological examination, placentas were fixed in formalin, weighed, examined grossly, and sectioned/stained. Routine sections were taken for histopathologic diagnoses per the Amsterdam Consensus Criteria recommendations.

### RNA extraction

Placental tissue was placed in Trizol (100 uL/10 mg) and homogenized using a TissueTearor. The resulting suspension was then centrifuged at 12,000 *x* g for 10 min after which the pellet was discarded. Chloroform was added to supernatant at a ratio of 100 uL chloroform/50 mg tissue. Tubes were shaken vigorously for 15 seconds, allowed to stand at room temperature for 10 min, and then centrifuged at 12,000 *x* g for 15 min. The aqueous phase was collected and the remainder of the RNA extraction procedure was performed using an RNeasy Mini Kit with on-column DNase I treatment (Qiagen) according to manufacturer instructions. RNA quantity and purity were assessed using a NanoDrop 2000 Spectrophotometer (ThermoFisher Scientific).

### cDNA synthesis and qPCR

cDNA synthesis was performed using the iScript^TM^ cDNA Synthesis Kit (Bio-Rad) according to manufacturer instructions. Briefly, 800ng of RNA was mixed with iScript Reaction Mix, iScript Reverse Transcriptase, and nuclease-free water. No template control and No RT controls were prepared. All samples were primed at 25°C for 5 min, heated to 46°C for 20 min, and RT was inactivated at 95°C for 1 min using a MiniAmpTM Plus Thermal Cycler (ThermoFisher Scientific). Quantitative real-time polymerase chain reaction (qPCR) was then performed using Taqman gene expression assays on a QuantStudio 5 Real-Time PCR System (ThermoFisher Scientific). Gene expression assays used are listed in Supplemental Table 4. Gene expression was normalized to the placental reference gene Tyrosine 3-Monooxygenase/Tryptophan 5-Monooxygenase Activation Protein Zeta (*YWHAZ*) and expressed relative to female Covid-19 negative samples to yield a relative quantity value (2^-ddCt^). All genes of interest were probed using gene expression assays with a dye of FAM-MGB, and *YWHAZ* was assessed using gene expression assays with a dye of VIC_PL. Two separate biopsies were each run in technical duplicate, and the average of biological replicates were then used to obtain final reported 2^-ddCt^ values.

### Protein extraction and quantification

Placental tissue was placed in RIPA Lysis and Extraction Buffer (ThermoFisher Scientific #89901) containing Halt Protease and Phosphatase Inhibitor Cocktail (ThermoFisher Scientific #78443) and homogenized using a TissueTearor. The resulting mixture was centrifuged at 10,000 rpm for 10 min at 4 degrees C and supernatant was kept for protein quantification. Protein was quantified using Pierce BCA Protein Assay Kit (ThermoFisher Scientific #23225).

### Immunoblotting

50 ug of protein was prepared in Protein Sample Loading Buffer (Li-Cor Biosciences) with 5 mM dithiothreitol (DTT), boiled for 10 min at 95 degree C, loaded on Mini-PROTEAN 4-20% TGX Stain-Free Precast Gels (Bio-Rad), and run at 200V for 30 min, after which they were transferred onto low fluorescence PVDF membrane using a Trans-Blot Turbo Transfer System (Bio-Rad). Blots were then washed 3x 10 min in TBS (20x stock from ThermoFisher diluted 1:20 in H2O) followed by imaging of Stain Free total protein with a ChemiDoc MP System (Bio-Rad). After stain free imaging, blots were washed 3x 10 min in TBS, followed by blocking in Intercept (TBS) Blocking Buffer (Li-Cor Biosciences) for 1 hr. Primary antibodies were then incubated in Intercept T20 (TBS) Antibody Diluent (Li-Cor Biosciences) overnight at 4°C. The next morning, membranes were washed 6x 10 min in TBS-T (20x stock from ThermoFisher was diluted 1:20 in H2O), followed by incubation with secondary antibodies at 1:15,000 dilution and hFAB Rhodamine Anti-Tubulin (Bio-Rad) at 1:4000 dilution for 1 hr in Intercept T20 (TBS) Antibody Diluent (Li-Cor Biosciences). Finally, blots were washed 6x 10 min in TBS-T, after which they briefly rinsed in and then placed in TBS. Fluorescence was then imaged with a ChemiDoc MP System (Bio-Rad) and quantified using Image Lab Software (Bio-Rad). Data is presented as volume of bands for protein of interest relative to the volume of bands contained in the total protein stain-free image.

### Immunofluorescence

For co-labeling immunofluoresence (IF) experiments, placenta tissue sections were rehydrated in an alcohol series after deparaffinization in xylene. Antigen retrieval was performed by boiling in 10 mM sodium citrate (pH 6.0) for 30min and cooled at room temperature before blocking for 15min with background sniper (Biocare medical). Samples were then incubated in primary antibodies diluted in 5% bovine serum albumin (BSA) for 1.5h at room temperature (Placental Alkaline Phosphatase (PLAP – ab212383) - 1:1000, Neonatal Fc Receptor (FcRn – ab193148) – 1:100, CD16 (CD16 – Leica NCL-L-CD16) – 1:100), CD32 (R&D AF1330) – 10ug/ml, CD64 (Origene TA506331)-1:100. The slides were washed in PBS Tween 0.1% and incubated in fluorescently conjugated secondary antibodies (1:400 in 5% BSA – Goat anti-Mouse IgG2a (A-21133), Goat anti-mouse IgG2b (A-21141), Goat anti-rabbit (ab150080), anti-mouse Igg1-(AF647 biolegend), donkey anti goat (AA1056)). Finally, the slides were treated with Vector True View to eliminate red blood cells and background (Vector True View - SP8400-15 - Vector Laboratories), treated with DAPI (D13060) and cover-slipped with vectashield mounting medium (Vector Laboratories -H-1000).

CellProfiler software (PMID:29969450) was used to quantify CD16/FcRn, CD32/FcRn, and CD64/FcRn colocalization as well as their respective intensities. Briefly, RGB pictures were converted to gray and placental vilii selected to avoid measuring background. Following background removal, the pictures were filtered using the function CorrectIlluminationCalculate and aligned to evaluate colocalization. Finally, CD16, CD32 or CD64 and FcRn intensities in the placental vilii previously delimitated to quantify colocalization.

CellProlifer software was similarly used to quantify immunohistochemistry for ACE2 and CD163. Following isolation of each dye compound (DAB and Hematoxylin), the placental vilii were manually isolated and the mean intensity of DAB signal quantified.

### Multivariate Analyses

All multivariate analyses were performed using R (version 4.0.0). Multivariate analyses included qPCR and Luminex antibody measurements. Missing data measurements were imputed using the k-nearest neighbor algorithm. Features were then centered and scaled to unit variance. Orthogonal partial least square discrimination analysis (O-PLSDA) was performed to classify data based on relative antibody transfer. Prior to building the O-PLSDA models, LASSO feature regularization and variable selection was performed, resulting in a reduced set of features that were consistently included in at least 80 out of 100 LASSO models. O-PLSDA models were built using the R ‘ropls’ Bioconductor package (orthI =1; PredI=1). Models were validated by evaluating the mean accuracy score after 100 trials of 5-fold CV. Accuracy scores for analogous models with permutated labels and random features were also reported for comparison.

### Statistical analyses

Statistical tests used and statistical significance are reported in the Figure Legends. Statistical analyses were performed using GraphPad Prism 9 and R (version 4.0.0). Significant differences in participant characteristics were evaluated using Kruskal-Wallis tests for continuous variables and chi-square or Fisher’s exact tests for categorical variables. Gene expression measurements, immunoblot measurements, and immunohistochemistry staining comparing maternal SARS-CoV-2 exposure status and fetal sex were evaluated by 2-way analysis of variance (2-way ANOVA) with Bonferroni’s post-hoc analysis. Kruskal-Wallis test followed by Dunn’s post-hoc test was used to determine significant differences between transfer ratios. Wilcoxon matched-pairs signed rank test was performed to determine significant differences between paired maternal-cord blood samples. All linear regression models were built in R (version 4.0.0) using the ‘stats’ package. False discovery rate (FDR) multiple comparisons correction was performed for Supplemental Fig. 6 using a corrected p-value of 0.05 as the cutoff for significance.

## SUPPLEMENTARY FIGURES AND TABLES

**Supplementary Figure 1.**
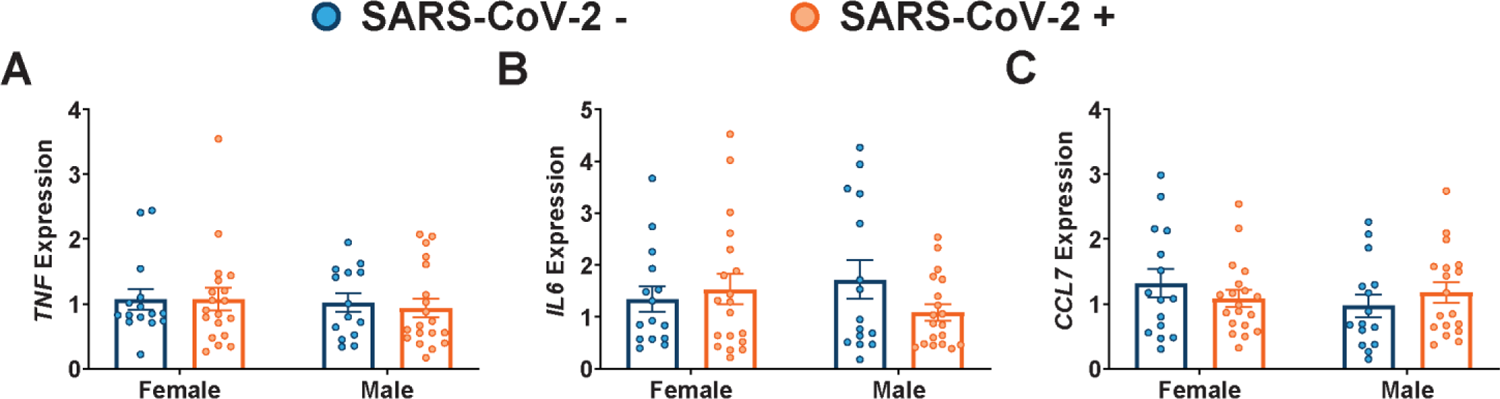
Maternal SARS-CoV-2 infection does not impact placental expression of *TNF, IL6,* or *CCL7*. **(A-C)** qPCR analyses of fetal male or fetal female expression of (A) *TNF,* (B) *IL6,* and (C) *CCL7* in placental biopsies from SARS-CoV-2 negative (blue) or SARS-CoV-2 positive (orange) pregnancies. Two-way ANOVA followed by post-hoc analyses were performed to determine significance. Expression levels shown are relative to reference gene *YWHAZ*. * p<0.05, ** p < 0.01.

**Supplementary Figure 2.**
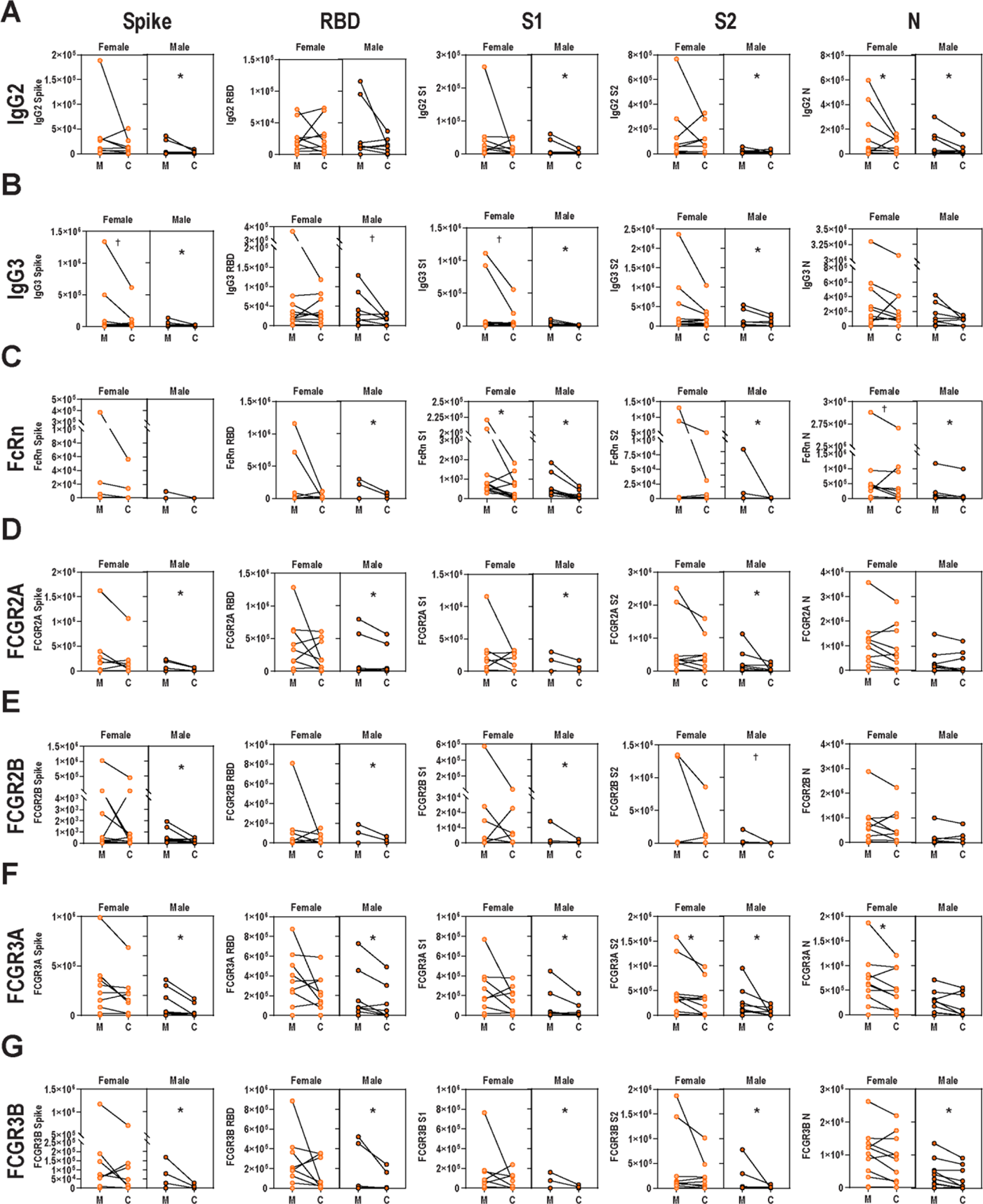
SARS-CoV-2 infected mothers with male fetuses demonstrate reduced placental transfer of SARS-CoV-2 antibodies compared to those with female fetuses. Dot plots showing relative Spike-, RBD-, S1-, S2-, and N-specific maternal blood (M) and cord blood (C) titers of (A) IgG2, (B), IgG3, (C) FcRn, (D) FCGR2A, (E) FCGR2B, (F) FCGR3A, and (G) FCGR3B. Females are shown in light orange while males are shown in dark orange with black border. Wilcoxon matched pairs signed rank test was performed to determine significance. *p<0.05, **p<0.01, ***p<0.0001.

**Supplementary Figure 3.**
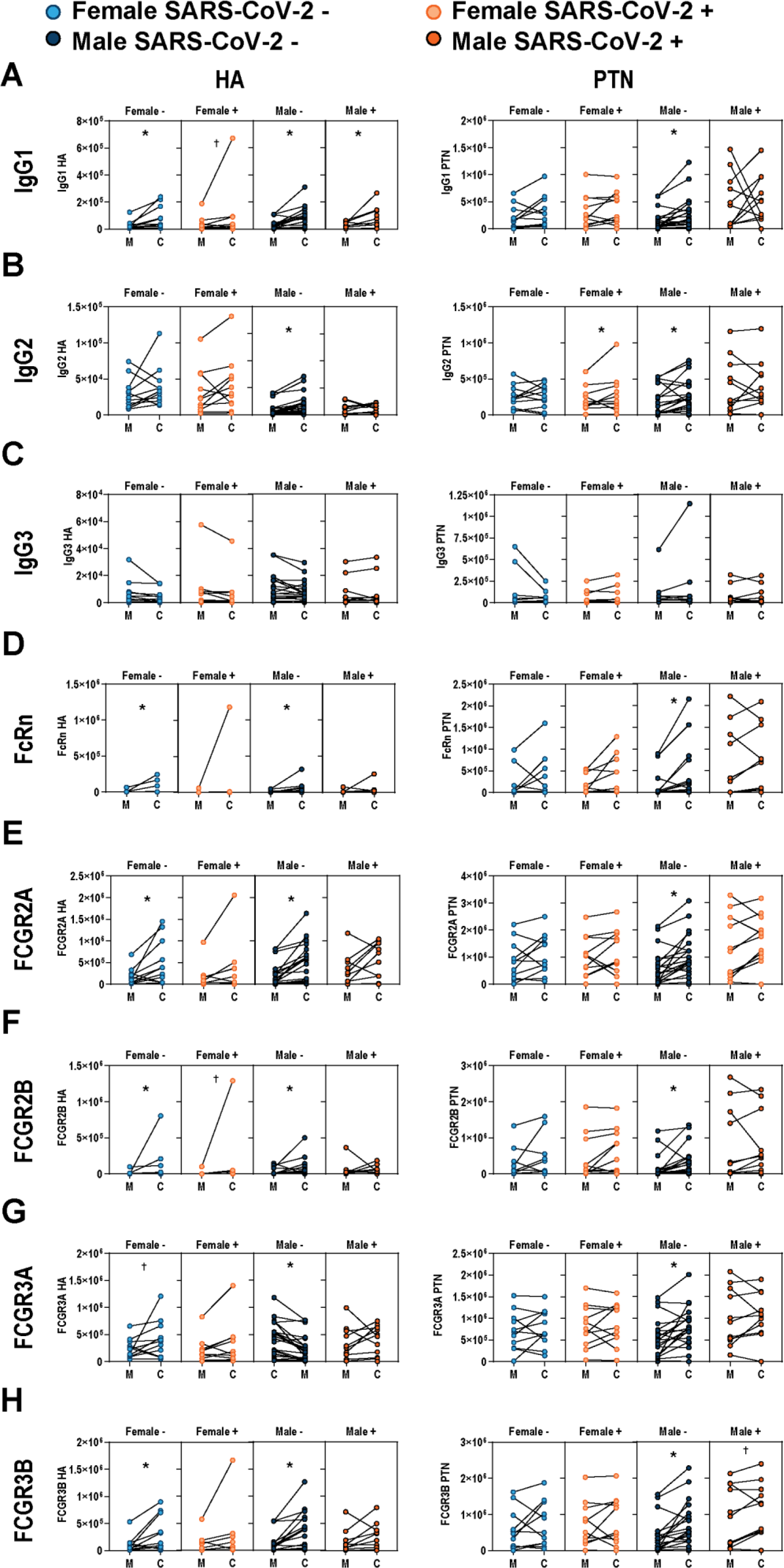
Maternal and cord blood titers of HA and PTN in SARS-CoV-2 infected and non-infected mothers. Dot plots showing relative hemagglutinin (HA)- and pertussis (PTN)-specific maternal blood (M) and cord blood (C) titers of (A) IgG1, (B) IgG2, (C) IgG3, (D) FcRn, (E) FCGR2A, (F) FCGR2B, (G) FCGR3A, and (H) FCGR3B. SARS-CoV-2 - females are shown in light blue, SARS-CoV-2 + females are shown in light orange, SARS-CoV-2 – males are shown in dark blue, and SARS-CoV-2 + males are shown in dark orange. Y-axis units for all plots are PBS-corrected mean fluorescence intensity (MFI). Wilcoxon matched-pairs signed rank test was performed to determine significance. * p < 0.05, † p<0.10.

**Supplementary Figure 4.**
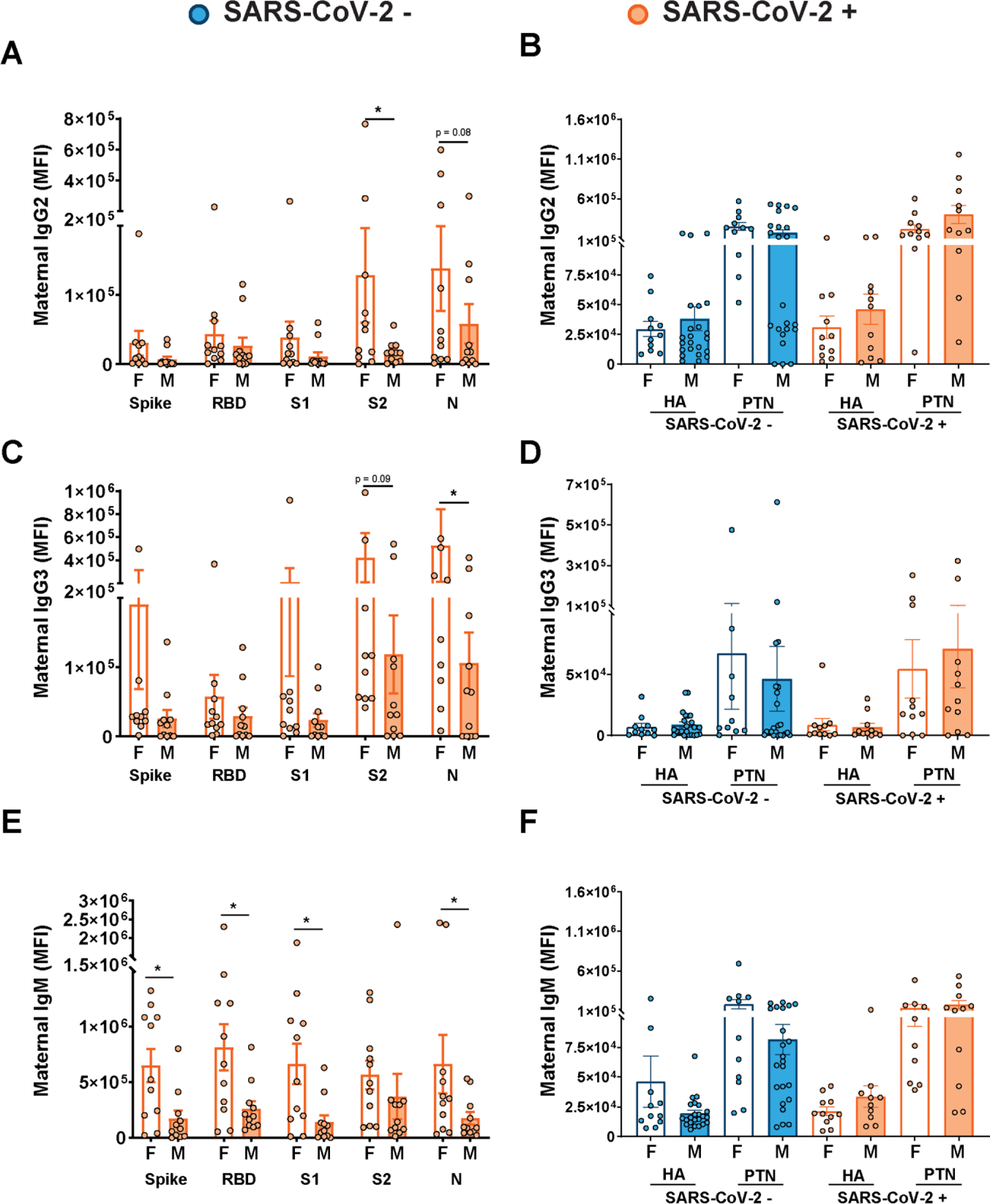
SARS-CoV-2 infected mothers with male fetuses have lower plasma levels of SARS-CoV-2-specific antibodies. **A.** Plots showing maternal Spike-, RBD-, S1-, S2-, and N-specific maternal blood IgG2 levels. Female neonates of mothers with SARS-CoV-2 are shown as white bars with an orange border while male neonates are shown as orange shaded bars with orange border. Two-way ANOVA followed by post-hoc analyses were performed to determine significance. There was a main effect of fetal/neonatal sex on maternal IgG1 levels (p = 0.003). *p < 0.05. **B.** Plots showing levels of IgG2 against HA and PTN in maternal plasma from either SARS-CoV-2 negative or SARS-CoV-2 positive mothers. Females born to SARS-CoV-2 – mothers are shown as white bars with blue border, females born to SARS-CoV-2 + mothers are shown as white bars with orange border. Males born to SARS-CoV-2 – mothers are shown as shaded blue bars with blue border, and males born to SARS-CoV-2 + mothers are shown as orange shaded bars with orange border. Two-way ANOVA revealed no significant effect of fetal sex. **C.** Plots showing maternal Spike-, RBD-, S1-, S2-, and N-specific maternal blood IgG3 levels. Two-way ANOVA followed by post-hoc analyses were performed to determine significance. There was a main effect of fetal sex on maternal IgG3 levels (p = 0.011). *p < 0.05. **D.** Plots showing levels of IgG3 against HA and PTN in maternal plasma from either SARS-CoV-2 negative or SARS-CoV-2 positive pregnancies. Two-way ANOVA revealed no significant effect of fetal sex. **E.** Plots showing maternal Spike-, RBD-, S1-, S2-, and N-specific maternal blood IgM3 levels. Two-way ANOVA followed by post-hoc analyses were performed to determine significance. There was a main effect of fetal sex on maternal IgM levels (p < 0.0001). *p < 0.05. **F.** Plots showing levels of IgM against HA and PTN in maternal plasma from either SARS-CoV-2 negative or SARS-CoV-2 positive pregnancies. Two-way ANOVA revealed no significant effect of fetal sex.

**Supplementary Figure 5.**
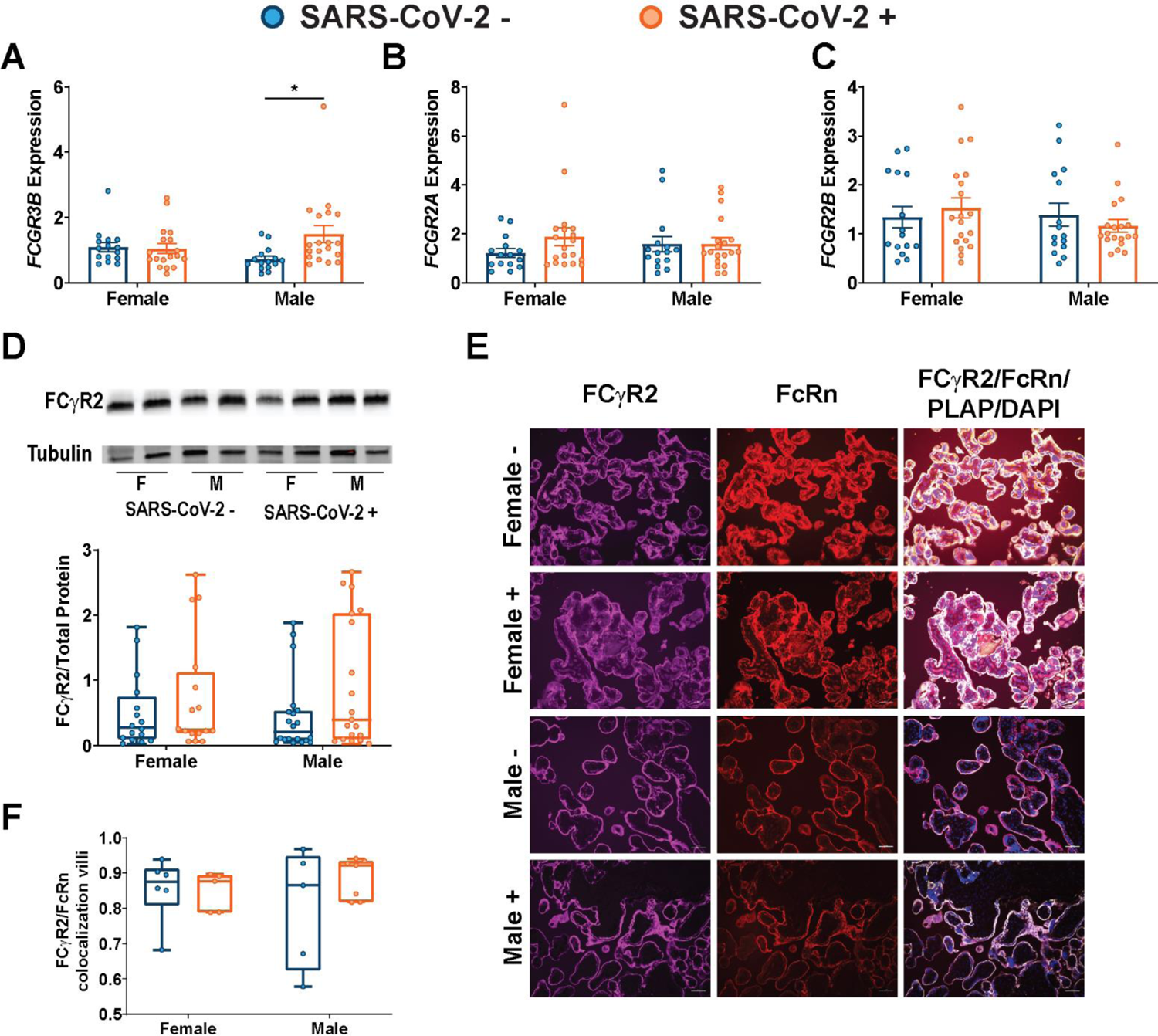
No effect of maternal SARS-CoV-2 infection on expression or localization of FCγR2. **(A-C)** qPCR analyses of fetal male or fetal female expression of (**A**) *FCGR3B,* (**B**) *FCGR2A,* and (**C**) *FCGR2B* in placental biopsies from SARS-CoV-2 negative (blue) or SARS-CoV-2 positive (orange) pregnancies. Expression levels shown are relative to reference gene *YWHAZ*. **(D)** Representative immunoblots and quantification of FCγR2 in female or male placental biopsies from mothers testing negative (blue) or positive (orange) for SARS-CoV-2. **(E)** Placental tissue sections from SARS-CoV-2 + and SARS-CoV-2 – mothers were stained for FCγR2 (purple), FcRn (red), and placental alkaline phosphatase (PLAP, green), a trophoblast marker, and DAPI (blue). **(F)** Box-and-whisker plots showing FCγR2/FcRn co-localization in placental villi. Two-way ANOVA followed by post-hoc analyses were performed to determine significance. * p<0.05.

**Supplementary Figure 6.**
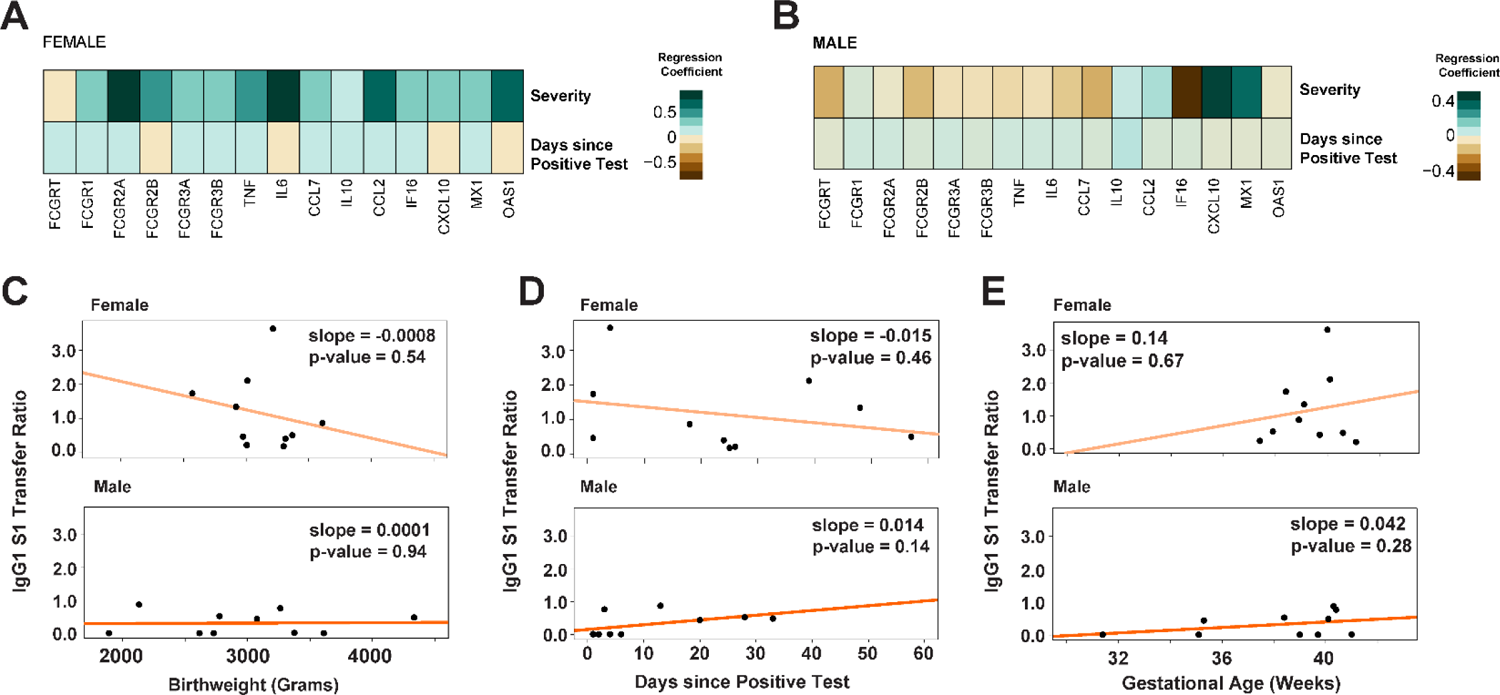
Effect of disease severity, time since infection, neonatal birthweight, and gestational age on gene expression and antibody transfer. **(A)** Heatmap depicting pairwise linear regressions between maternal disease severity (top), or time since positive test (bottom), and expression of each individual ISG or FcR for female placentas. None of the regression models had statistical significance after FDR multiple comparisons correction using a corrected p-value of 0.05 as the cutoff. **(B)** Heatmap depicting pairwise linear regressions between maternal disease severity (top), or time since positive test (bottom), and expression of each individual ISG or FcR for male placentas. None of the regression models had statistical significance after FDR multiple comparisons correction using a corrected p-value of 0.05 as the cutoff. **(C)** Linear regression models were performed on female (top) and male (bottom) neonates separately to examine the effect of birthweight on placental transfer of S1-specific IgG1 (no significant effect). **(D)** Linear regression models were performed on pregnancies with a female fetus (top) and male fetus (bottom) separately to examine the effect of days since maternal infection on transplacental transfer of S1-specific IgG1 (no significant effect). **(E)** Linear regression models were performed on female (top) and male (bottom) neonates separately to examine the effect of gestational age at delivery on placental transfer of S1-specific IgG1 (no significant effect).

**Supplementary Table 1.** Demographic and clinical characteristics of participants providing maternal and cord blood by fetal sex and maternal SARS-CoV-2 status. Abbreviations: DM/GDM, diabetes mellitus/gestational diabetes mellitus; BMI, body mass index; GA, gestational age; N/A, not applicable. ^a^SARS-CoV-2 positive and negative status determined by nasopharyngeal RT PCR at time of sample collection. If a participant was SARS-CoV-2 positive at any time in pregnancy she was included in “SARS-CoV-2 positive” category. ^b^ Significant differences between groups were determined using chi-square test for categorical variables, and Kruskal-Wallis test for continuous variables presented as median [IQR]. ^c^ Severity determinations were made based on published criteria from the Society for Maternal-Fetal Medicine and the National Institutes of Health.

|  | All (55) | Female |  | Male |  | *P*<sup>b</sup> |
| --- | --- | --- | --- | --- | --- | --- |
|  |  | SARS-CoV-2 negative (11) | SARS-CoV-2 positive (11)<sup>a</sup> | SARS-CoV-2 negative (22) | SARS-CoV-2 positive (11)<sup>a</sup> |  |
| Maternal Age, years | 33 [29-38] | 34 [32-38] | 37 [28-40] | 32 [29-37] | 31 [25-36] | 0.43 |
| Parity, *n* | 1 [0-2] | 1 [0-2] | 1 [0-1] | 1 [0-2] | 1 [0-2] | 0.95 |
| Race, *n* (%) |  |  |  |  |  | 0.03 |
| *White* | 36 (65) | 8 (73) | 5 (45) | 18 (82) | 5 (45) |  |
| *Black* | 4 (7) | 0 (0) | 3 (27) | 0 (0) | 1 (9) |  |
| *Asian* | 2 (4) | 0 (0) | 0 (0) | 2 (9) | 0 (0) |  |
| *Other* | 8 (15) | 3 (27) | 2 (18) | 1 (5) | 2 (18) |  |
| *Not Reported* | 5 (9) | 0 (0) | 1 (9) | 1 (5) | 3 (27) |  |
| Ethnicity, *n* (%) |  |  |  |  |  | 0.01 |
| *Hispanic* | 19 (35) | 3 (27) | 4 (36) | 3 (14) | 9 (82) |  |
| *Non-Hispanic* | 33 (60) | 7 (64) | 6 (55) | 18 (82) | 2 (18) |  |
| *Not Reported* | 3 (5) | 1 (9) | 1 (9) | 1 (5) | 0 (0) |  |
| Chronic hypertension, *n* (%) | 1 (2) | 0 (0) | 0 (0) | 1 (5) | 0 (0) | 0.68 |
| DM/GDM, *n* (%) | 6 (11) | 1 (9) | 1 (9) | 3 (14) | 1 (9) | 0.96 |
| BMI ≥ 30 kg/m<sup>2</sup>, *n* (%) | 17 (31) | 2 (18) | 6 (55) | 5 (23) | 4 (36) | 0.21 |
| Pre-pregnancy BMI, kg/m<sup>2</sup> | 27.0 [21.6-32.0] | 27.0 [22.6-29.8] | 29.2 [25.9-36.3] | 22.1 [20.3-30.3] | 29.5 [22.5-32.1] | 0.16 |
| GA at delivery, weeks | 39 [38-40] | 39 [39-39] | 40 [38-41] | 39 [38-41] | 39 [35-40] | 0.86 |
| Any labor, *n* (%) | 40 (73) | 4 (36) | 10 (91) | 16 (73) | 10 (91) | 0.01 |
| Neonatal birthweight, *g* | 3300 [3005-3630] | 3295 [3060-3590] | 3010 [2920-3315] | 3533 [3254-3784] | 3075 [2615-3610] | 0.07 |
| Multiple gestation, *n* | 1 | 0 | 0 | 1 | 0 | N/A |
| GA at positive SARS-CoV-2 test, weeks | 36.3 [32.4-38.8] | N/A | 36.3 [32.4] | N/A | 35.4 [32.4-39.6] | 0.81 |
| COVID-19 disease severity at diagnosis<sup>c</sup>, *n* |  |  |  |  |  | 0.77 |
| *Asymptomatic* | 5 (23) | N/A | 2 (18) | N/A | 3 (27) |  |
| *Mild/Moderate* | 14 (64) | N/A | 7 (64) | N/A | 7 (64) |  |
| *Severe/Critical* | 3 (14) | N/A | 2 (18) | N/A | 1 (9) |  |
| Time between SARS-CoV-2 symptom onset and delivery, days | 29 [9-44] | N/A | 33 [26-58] | N/A | 29 [9-44] | 0.08 |

**Supplementary Table 2.** **2-way ANOVA analysis of inflammatory cytokine and interferon stimulated gene expression.** Two-way ANOVA followed by Bonferroni’s post-hoc analyses (when interaction term was significant) were performed to determine significance. All main and interaction effects for genes of interest are represented for both male and female placentas in addition to all post-hoc analyses performed. Significant effects are indicated by bolded statistics. N/A: not applicable, indicates post-hoc testing not performed due to lack of significant interaction term.

| Gene | Maternal SARS-CoV-2 Status | Fetal Sex | Interaction | Bonferroni Female Neg:Pos | Bonferroni Male Neg:Pos | Bonferroni Neg Female:Male | Bonferroni Pos Female:Male |
| --- | --- | --- | --- | --- | --- | --- | --- |
| *TNF* | $F_{(1,63)} = 0.061, p = 0.81$ | $F_{(1,63)} = 0.31, p = 0.58$ | $F_{(1,63)} = 0.076, p = 0.78$ | n/a | n/a | n/a | n/a |
| *IL6* | $F_{(1,64)} = 0.66, p = 0.42$ | $F_{(1,64)} = 0.017, p = 0.90$ | $F_{(1,64)} = 2.3, p = 0.13$ | n/a | n/a | n/a | n/a |
| *CCL7* | $F_{(1,63)} = 0.0075, p = 0.93$ | $F_{(1,63)} = 0.57, p = 0.45$ | $F_{(1,63)} = 1.71, p = 0.20$ | n/a | n/a | n/a | n/a |
| *IL10* | $F_{(1,63)} = 3.58, p = 0.063$ | $F_{(1,63)} = 0.19, p = 0.66$ | **$F_{(1,63)} = 7.72, p = 0.007$** | $p > 0.9999$ | **$p = 0.003$** | $p = 0.068$ | $p = 0.17$ |
| *CCL2* | **$F_{(1,64)} = 4.69, p = 0.03$** | $F_{(1,64)} = 0.85, p = 0.36$ | $F_{(1,64)} = 1.25, p = 0.27$ | n/a | n/a | n/a | n/a |
| *IFI6* | $F_{(1,64)} = 0.25, p = 0.62$ | $F_{(1,64)} = 0.17, p = 0.68$ | **$F_{(1,64)} = 7.74, p = 0.007$** | $p = 0.22$ | **$p = 0.047$** | $p = 0.24$ | **$p = 0.038$** |
| *CXCL10* | $F_{(1,62)} = 3.049, p = 0.086$ | $F_{(1,62)} = 0.17, p = 0.68$ | **$F_{(1,62)} = 12.36, p = 0.0008$** | $p = 0.42$ | **$p = 0.001$** | **$p = 0.022$** | **$p = 0.045$** |
| *MX1* | **$F_{(1,63)} = 4.46, p = 0.039$** | $F_{(1,63)} = 0.12, p = 0.74$ | $F_{(1,63)} = 1.038, p = 0.31$ | n/a | n/a | n/a | n/a |
| *OAS1* | $F_{(1,63)} = 0.74, p = 0.39$ | $F_{(1,63)} = 0.039, p = 0.85$ | **$F_{(1,63)} = 5.99, p = 0.017$** | $p = 0.54$ | **$p = 0.044$** | $p = 0.27$ | $p = 0.11$ |

**Supplementary Table 3.**
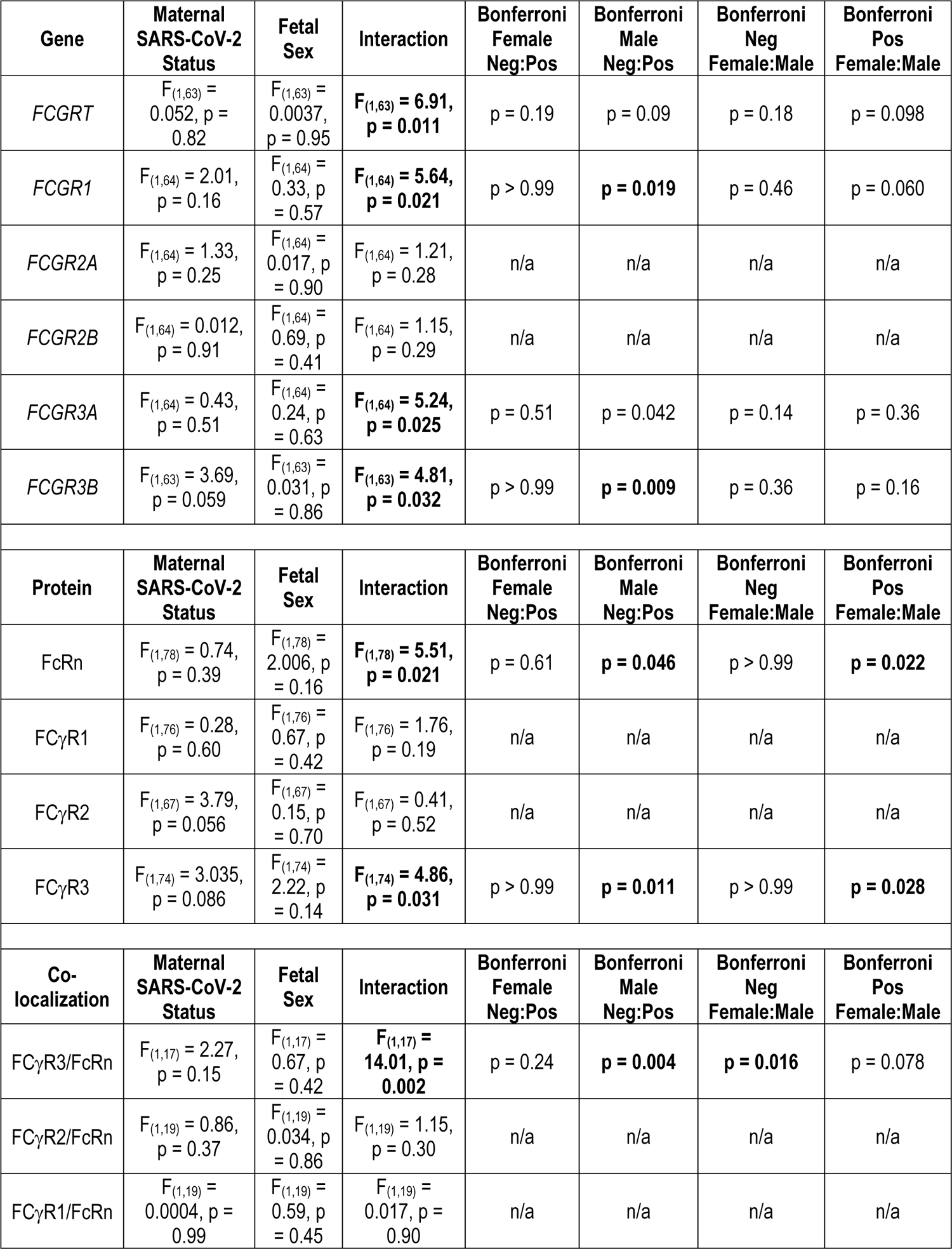
**2-way ANOVA analysis of Fc receptor gene expression, immunoblots, and immunohistochemistry.** Two-way ANOVA followed by Bonferroni’s post-hoc analyses (when interaction term was significant) were performed to determine significance. All main and interaction effects for genes and proteins of interest are represented for both male and female placentas in addition to all post-hoc analyses performed. Significant effects are indicated by bolded statistics. N/A: not applicable, indicates post-hoc testing not performed due to lack of significant interaction term.

| Gene | Maternal SARS-CoV-2 Status | Fetal Sex | Interaction | Bonferroni Female Neg:Pos | Bonferroni Male Neg:Pos | Bonferroni Neg Female:Male | Bonferroni Pos Female:Male |
| --- | --- | --- | --- | --- | --- | --- | --- |
| *FCGRT* | $F_{(1,63)} = 0.052, p = 0.82$ | $F_{(1,63)} = 0.0037, p = 0.95$ | $F_{(1,63)} = 6.91, p = 0.011$ | $p = 0.19$ | $p = 0.09$ | $p = 0.18$ | $p = 0.098$ |
| *FCGR1* | $F_{(1,64)} = 2.01, p = 0.16$ | $F_{(1,64)} = 0.33, p = 0.57$ | $F_{(1,64)} = 5.64, p = 0.021$ | $p > 0.99$ | $p = 0.019$ | $p = 0.46$ | $p = 0.060$ |
| *FCGR2A* | $F_{(1,64)} = 1.33, p = 0.25$ | $F_{(1,64)} = 0.017, p = 0.90$ | $F_{(1,64)} = 1.21, p = 0.28$ | n/a | n/a | n/a | n/a |
| *FCGR2B* | $F_{(1,64)} = 0.012, p = 0.91$ | $F_{(1,64)} = 0.69, p = 0.41$ | $F_{(1,64)} = 1.15, p = 0.29$ | n/a | n/a | n/a | n/a |
| *FCGR3A* | $F_{(1,64)} = 0.43, p = 0.51$ | $F_{(1,64)} = 0.24, p = 0.63$ | $F_{(1,64)} = 5.24, p = 0.025$ | $p = 0.51$ | $p = 0.042$ | $p = 0.14$ | $p = 0.36$ |
| *FCGR3B* | $F_{(1,63)} = 3.69, p = 0.059$ | $F_{(1,63)} = 0.031, p = 0.86$ | $F_{(1,63)} = 4.81, p = 0.032$ | $p > 0.99$ | $p = 0.009$ | $p = 0.36$ | $p = 0.16$ |
| Protein | Maternal SARS-CoV-2 Status | Fetal Sex | Interaction | Bonferroni Female Neg:Pos | Bonferroni Male Neg:Pos | Bonferroni Neg Female:Male | Bonferroni Pos Female:Male |
| FcRn | $F_{(1,78)} = 0.74, p = 0.39$ | $F_{(1,78)} = 2.006, p = 0.16$ | $F_{(1,78)} = 5.51, p = 0.021$ | $p = 0.61$ | $p = 0.046$ | $p > 0.99$ | $p = 0.022$ |
| FCγR1 | $F_{(1,76)} = 0.28, p = 0.60$ | $F_{(1,76)} = 0.67, p = 0.42$ | $F_{(1,76)} = 1.76, p = 0.19$ | n/a | n/a | n/a | n/a |
| FCγR2 | $F_{(1,67)} = 3.79, p = 0.056$ | $F_{(1,67)} = 0.15, p = 0.70$ | $F_{(1,67)} = 0.41, p = 0.52$ | n/a | n/a | n/a | n/a |
| FCγR3 | $F_{(1,74)} = 3.035, p = 0.086$ | $F_{(1,74)} = 2.22, p = 0.14$ | $F_{(1,74)} = 4.86, p = 0.031$ | $p > 0.99$ | $p = 0.011$ | $p > 0.99$ | $p = 0.028$ |
| Co-localization | Maternal SARS-CoV-2 Status | Fetal Sex | Interaction | Bonferroni Female Neg:Pos | Bonferroni Male Neg:Pos | Bonferroni Neg Female:Male | Bonferroni Pos Female:Male |
| FCγR3/FcRn | $F_{(1,17)} = 2.27, p = 0.15$ | $F_{(1,17)} = 0.67, p = 0.42$ | $F_{(1,17)} = 14.01, p = 0.002$ | $p = 0.24$ | $p = 0.004$ | $p = 0.016$ | $p = 0.078$ |
| FCγR2/FcRn | $F_{(1,19)} = 0.86, p = 0.37$ | $F_{(1,19)} = 0.034, p = 0.86$ | $F_{(1,19)} = 1.15, p = 0.30$ | n/a | n/a | n/a | n/a |
| FCγR1/FcRn | $F_{(1,19)} = 0.0004, p = 0.99$ | $F_{(1,19)} = 0.59, p = 0.45$ | $F_{(1,19)} = 0.017, p = 0.90$ | n/a | n/a | n/a | n/a |

**Supplementary Table 4.** Taqman gene expression assays used for RTqPCR. ThermoFisher Scientific Assay ID, Gene symbol, and color dye for each gene expression assay used.

| **Assay ID** | **Gene Symbol** | **Dye Label** |
| --- | --- | --- |
| Hs00237047_m1 | *YWHAZ* | VIC_PL |
| Hs00175415_m1 | *FCGRT* | FAM-MGB |
| Hs00174081_m1 | *FCGR1A* | FAM-MGB |
| Hs00234969_m1 | *FCGR2A* | FAM-MGB |
| Hs00269610_m1 | *FCGR2B* | FAM-MGB |
| Hs02388314_m1 | *FCGR3A* | FAM-MGB |
| Hs04334165_m1 | *FCGR3B* | FAM-MGB |
| Hs00234140_m1 | *CCL2* | FAM-MGB |
| Hs00174131_m1 | *IL6* | FAM-MGB |
| Hs00174128_m1 | *TNF* | FAM-MGB |
| Hs00171147_m1 | *CCL7* | FAM-MGB |
| Hs00242571_m1 | *IFI6* | FAM-MGB |
| Hs00171042_m1 | *CXCL10* | FAM-MGB |
| Hs00895608_m1 | *MX1* | FAM-MGB |
| Hs00961622_m1 | *IL10* | FAM-MGB |
| Hs00242943_m1 | *OAS1* | FAM-MGB |

